# Small-molecule targeting MuRF1 protects against denervation-induced diaphragmatic dysfunction: Underlying molecular mechanisms

**DOI:** 10.1101/2025.07.03.662973

**Authors:** Fernando Ribeiro, Paulo R. Jannig, Siegfried Labeit, Anselmo S. Moriscot

## Abstract

**Background:** Mechanical inactivity rapidly induces diaphragm muscle fibers’ contractile dysfunction and atrophy. Diaphragm weakness can impair respiratory function, quality of life, and increase risks of morbidity and mortality. Muscle RING-finger protein-1 (MuRF1) expression is upregulated during denervation and muscle inactivity, and is known to target key muscle proteins for degradation. We previously reported that the small-molecule targeting MuRF1 (MyoMed-205) protects against diaphragm contractile dysfunction and atrophy after 12 hours of unilateral diaphragm denervation (UDD) in rats. In this study, we investigated the mechanisms by which MyoMed-205 protects the diaphragm structure and function during early UDD in rats.

**Methods:** Male Wistar rats were subjected to unilateral diaphragm denervation (UDD) for 12 hours. Immediately after UDD, rats received either a placebo (vehicle) or small-molecule targeting MuRF1 (MyoMed-205, 50 mg/kg bw), and outcomes were compared with Sham-operated controls. Diaphragm was used for histological, morphometric, transcriptomic (RNA-seq), and protein content (Western Blot) analysis.

**Results:** UDD induced diaphragm slow-(type I: p = 0.03) and fast-twitch (type IIa: p = 0.04; type IIb/x: p = 0.02) fibers atrophy after 12 hours, which was prevented by MyoMed-205 (p < 0.05). Mechanistically, UDD perturbed mechanisms involved with myofiber ultrastructure and contractility, mitochondrial function, proteolysis, and tissue remodeling in the diaphragm. MyoMed-205 enhanced the activation of mechanisms required for sarcomere integrity, calcium handling, antioxidant defense, chaperone-mediated unfolded protein response, and muscle growth. MyoMed-205 also mitigated intramuscular fat deposition and pro-fibrotic responses triggered by UDD.

**Conclusion:** Small-molecule targeting MuRF1 (MyoMed-205) protects against diaphragm muscle contractile dysfunction and atrophy after 12 hours of UDD. Herein, we demonstrate that this protective effect involved augmented activation of signaling pathways controlling muscle structure and function, chaperone-mediated unfolded protein, and muscle growth, while mitigating intramuscular fat deposition and pro-fibrotic responses triggered by UDD at the transcriptional and/or protein level.

## Introduction

Breathing is essential for life. This key biological process is regulated by a complex neuromuscular network generating intrathoracic pressures for lung ventilation and gas exchange. The diaphragm functions as the primary inspiratory pump while contributing to other physiological processes (e.g., visceral organization, coughing, and venous hemodynamics). Due to unique evolutionary adaptations and high lifelong contractile activity, the diaphragm might respond distinctly to mechanical inactivity compared to other skeletal muscles [1]. Clinical conditions such as phrenic nerve injury (e.g., thoracic surgery, trauma) and prolonged mechanical ventilation cause profound diaphragm structural and functional changes. Resulting diaphragm muscle weakness and wasting can impair respiratory capacity, exercise tolerance, quality of life, and increase mortality [2].

Understanding how mechanical activity affects diaphragm muscle structure and function has special clinical relevance [for review, see [3,4]. The unilateral diaphragm denervation (UDD) model has been used to investigate the diaphragm adaptations to short- and long-term periods of absent innervation and contractile function. Even short periods (e.g., < 24 hours) of inactivity following UDD significantly decrease diaphragm force-generation capacity [5,6]. Within 24 hours, contractile function loss can accompany slow- and fast-twitch fibers atrophy [6,7], though other findings suggest that fiber size may be less affected initially, especially in slow-twitch fibers [5]. Adaptive responses triggered by UDD include diaphragm ultrastructural changes, myofibrillar protein loss, altered protein balance, increased activation of proteolytic pathways, apoptosis, and muscle tissue remodeling [6,8–13].

Since its discovery in the early 2000s, Muscle RING-finger protein 1 (MuRF1) has been recognized as a key regulator of skeletal muscle mass and function [14,15]. This muscle-specific E3-ligase targets protein clusters controlling muscle contraction, Ca2+ handling, energy metabolism, immune response, and protein breakdown [16–18]. Increased MuRF1 expression/activity occurs in several stress states (e.g., immobilization, denervation, corticosteroids, heart failure, cancer cachexia, aging) and is associated with muscle wasting and weakness, while MuRF1 deletion improves muscle mass and function [19]. Recent advances in small-molecule MuRF1 inhibitors have shown therapeutic potential across multiple conditions, attenuating muscle wasting in heart failure [22–24], cancer cachexia [25], corticosteroid-induced myopathies [22], and diabetes [26]. Notably, MuRF1 deletion or inhibitor protects against early diaphragmatic dysfunction in mechanically ventilated rodents [20,21], and our group previously demonstrated that small-molecule MuRF1 targeting (MyoMed-205) prevents early diaphragm contractile dysfunction and atrophy in rats subjected to UDD [6], suggesting MuRF1 as a promising therapeutic target for counteracting denervation-induced diaphragm muscle weakness and wasting.

This study aimed to investigate the cellular and molecular mechanisms underlying the small-molecule targeting (MyoMed205) mediated protection against diaphragm dysfunction and atrophy in rats subjected to 12 hours of UDD. We hypothesized that 1) MuRF1 plays a key role in denervation-induced diaphragmatic dysfunction and atrophy, and 2) MyoMed-205 differentially regulates signaling pathways controlling diaphragm muscle contractile function, mass, and tissue remodeling during denervation stress.

## Material and Methods

### Animals

Wistar rats (Male, 2-3 months old) were used in this study. The animals were kept in standard cages under controlled environmental conditions (24 ± 1°C, 12 h/12 h light-dark cycle) with access to standard food and water *ad libitum*. This study was approved and followed the institutional guidelines for animal care and use for research (CEUA ICB USP #8728030320 and #5143091020).

### Experimental design

Twenty-four rats were randomly allocated into three experimental groups (*n* = 8): Sham 12h (Sham); UDD 12 hours + vehicle (DNV + VEH); and UDD 12 hours + MyoMed-205 (DNV + 205). After 12 hours, the animals were euthanized under anesthesia, and the denervated right diaphragm muscle (costal portion) was harvested, quickly snap-frozen in 2-methylbutane chilled in liquid nitrogen, and stored at −80°C until further analysis. Diaphragm samples used in this study originate from our previous work [6] (for further details, see supplementary information).

### MyoMed-205 formulation and delivery

A detailed description of MuRF1 inhibitor (MyoMed-205, Myomedix GmbH, Germany) formulation for the *in vivo* study is provided in the supplementary information. Briefly, the compound was freshly dissolved in an 8 mL vehicle solution (DMSO-PEG400-Saline 0.9%). The delivery of either vehicle (VEH) or MyoMed-205 (205) treatments was managed in four split doses (2 mL/dose) administered every 3 h interval over the 12-hour experimental period via the caudal vein. Treatments started immediately following the confirmation of unilateral diaphragm paralysis (for further details, see supplementary information).

### Unilateral diaphragm denervation (UDD)

Animals were initially sedated and anesthetized via intraperitoneal administration of acepromazine (2.5 mg/kg bw), followed after 30 minutes by the administration of ketamine (100 mg/kg bw) and xylazine (5 mg/kg bw). After reaching the surgical plane, the right phrenic nerve was exposed and transected at the rat’s lower neck region. The absence of right diaphragm dome contractile activity was used to confirm the denervation’s efficacy. Then, the surgical wound was sutured and managed with antiseptics. During the post-operative period, animals were allowed to recover from anesthesia in individual cages with *ad libitum* access to food chow and water. (for further details, see supplementary information).

### Diaphragm muscle fiber typing and size analysis

Cryosections (8 µm thick) from the right costal diaphragm muscle were prepared using a cryostat at −20°C (Leica CM1850, Germany). Muscle fiber typing (MyHC I, IIA, IIB/X) was performed via immunofluorescence assay (see antibodies in Table S1). Photomicrographs were acquired using Axio Scope A1 microscope (Zeiss, Germany). Diaphragm fibers’ type distribution and cross-sectional area (CSA) were assessed with ImageJ (Fiji, NIH, USA). Representative muscle fiber type distribution and CSA values were determined as the average from 300 measurements per sample of previously unanalyzed muscle fibers (for further details, see supplementary information).

### Oil Red O (ORO) Staining

Diaphragm muscle cryosections were stained with ORO to assess the intramuscular lipid content. Photomicrographs were acquired using Axio Scope A1 microscope (Zeiss, Germany). Representative number of positive ORO (ORO+) muscle cells and intracellular lipid content were determined based on measurements of 300 fibers per sample (for further details, see supplementary information).

### Transcriptomic profiling (RNA-seq)

Total RNA was extracted from the diaphragm using TRIzol (Life Technologies, Carlsbad, CA). RNA integrity was analyzed using Bioanalyzer 2100 (Agilent, Santa Clara, CA, USA) (Table S2). Four samples of each experimental group with the highest purity and integrity levels were selected for global gene expression analysis (Figure S1). Approximately ∼1 ug of sample total RNA was used for cDNA library preparation performed according to the Illumina mRNA stranded poly-A tail enrichment Kit manufacturer’s instructions (Illumina, San Diego, CA). RNA sequencing was carried out using the Illumina NextSeq 2000 (NextSeq 2x100pb, 20 million paired-end reads depth). These procedures were performed by NGS Soluções Genômicas (Piracicaba, Brazil).

For bioinformatic analysis, quality control of raw reads was performed using the FastQC toolkit (Babraham Bioinformatics). The reads were then aligned to the mRatBN7.2 rat genome using STAR. Feature count was performed using featureCounts, and differential expression analyses were generated in R using the DESeq2 package with adaptive log-fold change shrinkage estimator from the ashr package Pathway analysis was carried out by Gene Set Enrichment Analysis (GSEA) using a pre-ranked list of genes by log2Fold-Change and Molecular Signatures Database, and hallmark pathways gene sets for pathway analysis. Transcriptome-related raw files and data can be found in the Gene Expression Omnibus (GEO) repository (GSE305304) (for further details, see supplementary information).

### Western Blotting

Diaphragm samples were powdered in a liquid nitrogen-chilled mortar and homogenized in RIPA buffer containing protease and phosphatase inhibitors (ThermoFisher #78445). Homogenates were centrifuged and the collected and stored at −80°C until further analysis. Protein concentration was determined by the Bradford method with bovine serum albumin (BSA) as the standard. Protein was loaded onto polyacrylamide gel (SDS-PAGE), separated via electrophoresis (100V), and transferred to polyvinylidene difluoride (PVDF) membrane (Thermo Fisher Scientific, #88518, Rockford, IL, USA) using wet-transfer system (100 V, 350 mA, for 120 min). Ponceau red staining was used to assess protein transfer efficacy. Membranes were incubated with primary and secondary antibodies (Table S1). Membranes were incubated with ECL substrate (Immobilon Forte, Millipore #WBLUF0500) for 5 minutes before signal detection using C-digit blot scanner (Li-Cor, USA) for chemiluminescent detection, or with BCIP/NBT substrate (Sigma #B1911) for colorimetric detection. Alpha-tubulin was used as a loading control for data normalization, and densitometry analysis was performed using ImageStudio software (for further details, see supplementary information).

### Statistical Analysis

Data analysis was performed Excel, Prism and R. For the analysis of the transcriptomic data, refer to the respective section above and supplementary information. For the analysis of histological, morphometric, and protein expression levels, the Shapiro-Wilk test was used to assess data normality. One-way ANOVA followed by Tukey’s post-hoc test, or the Kruskal-Wallis followed by Dunn’s test, were used for multiple comparisons between groups, for parametric and non-parametric data, respectively. Data are presented as mean ± standard deviation and statistical significance was accepted as *p* < 0.05.

## Results

### Transcriptomic profiling reveals mechanisms underlying MyoMed-205-mediated protection against denervation-induced diaphragmatic dysfunction and atrophy

To investigate how denervation might impair diaphragm muscle structure and contractile function, as reported previously [6], we performed a global gene expression analysis of the rat’s diaphragm following 12 hours of UDD. Additionally, we compared the transcriptomes of MyoMed-205-treated animals with untreated animals to investigate the mechanisms underlying the MyoMed-205-mediated protective effects. For this purpose, animals were treated either with vehicle (DNV_VEH) or MyoMed-205 (DNV_205), and results were compared with Sham controls (Figure 1A).

**Figure 1.**
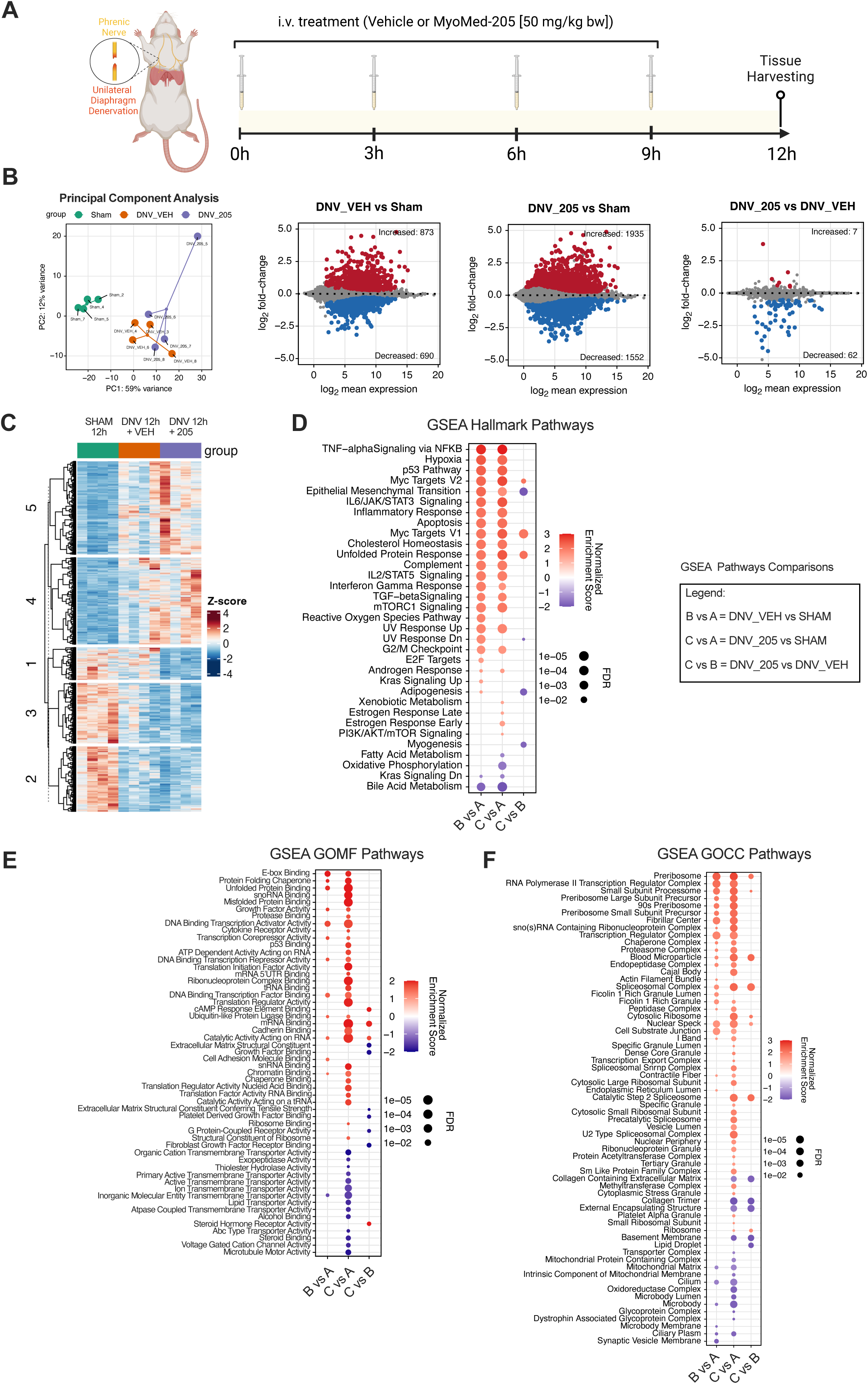
Transcriptomic profiling of the diaphragm following 12 h of unilateral diaphragm denervation (UDD) and identification of pathways responsive to MyoMed-205. (A) Schematic illustration of the experimental design used to investigate the effect of small-molecule targeting MuRF1 (MyoMed-205, 50 mg/kg bw, intravenous delivery) upon early UDD pathophysiology in rats. (B) Principal component analysis for RNA- seq of diaphragm muscle and identification of differentially expressed genes (DEGs, Padj < 0.05) across the distinct experimental conditions (*n* = 4 per group). Upregulated genes are represented in red, while downregulated genes are represented in blue. (C) Hierarchical clustered heat map highlights five clusters of DEGs identified in response to UDD and MyoMed-205 compared with the control. (D) Gene Set Enrichment Analysis (GSEA) reveals the hallmark pathways responsive to UDD and the impacts of MyoMed-205. Additionally, GSEA indicates the altered (E) molecular functions (GOMF) and (F) cellular compartments (GOCC) of the diaphragm muscle in response to UDD and MyoMed-205 compared with controls. Padj, adjusted p-value. FDR, false discovery rate.

Principal component analysis (PCA) demonstrated significant transcriptomic alterations in DNV_VEH compared with Sham. On the other hand, MyoMed-205 caused no evident changes in the clustering pattern compared with DNV_VEH (Figure 1B). DESeq2 analysis was performed to identify differentially expressed genes (DEGs) for each condition (Padj < 0.05). We detected 1563 DEGs in DNV_VEH (873 upregulated, 690 downregulated) and 3487 DEGs in DNV_205 (1935 upregulated, 1552 downregulated), compared with Sham. We found 69 DEGs particularly responsive to MyoMed-205 treatment compared with DNV_VEH (Figure 1B), of which 7 were upregulated and 62 downregulated (see Table 1). Hierarchical clustering analysis revealed five gene clusters regulated by UDD and MyoMed-205 (Figure 1C).

**Table 1.**
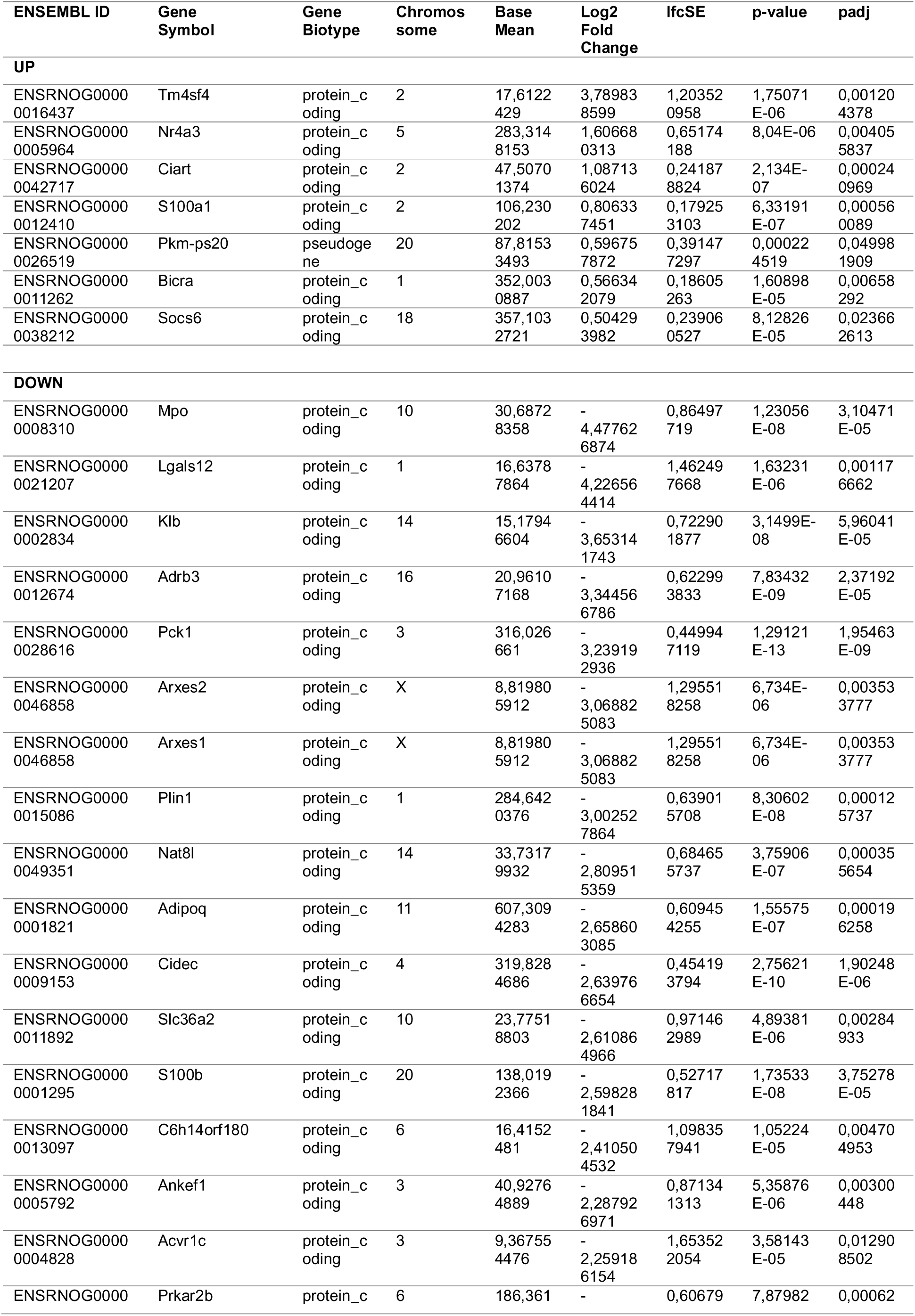
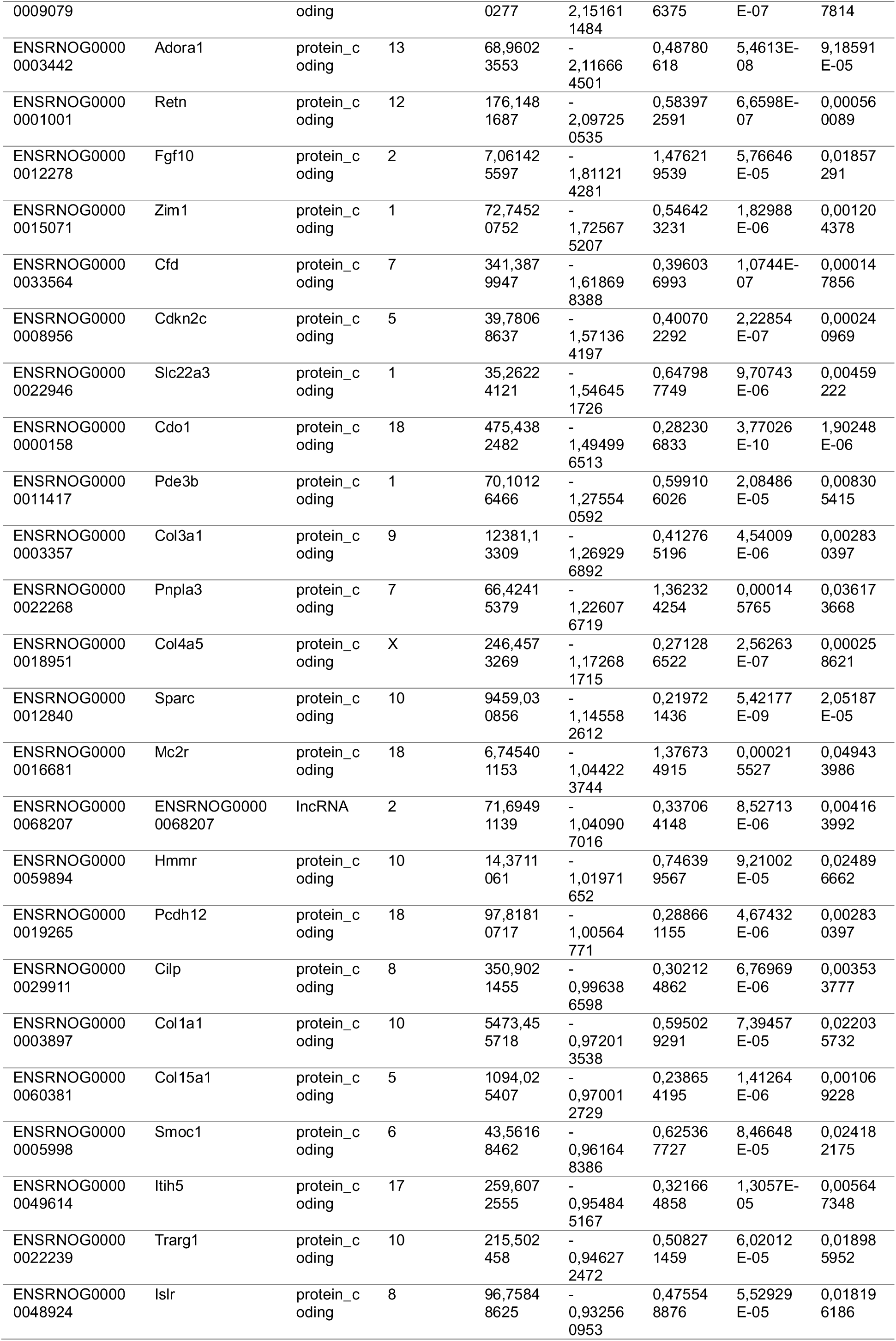
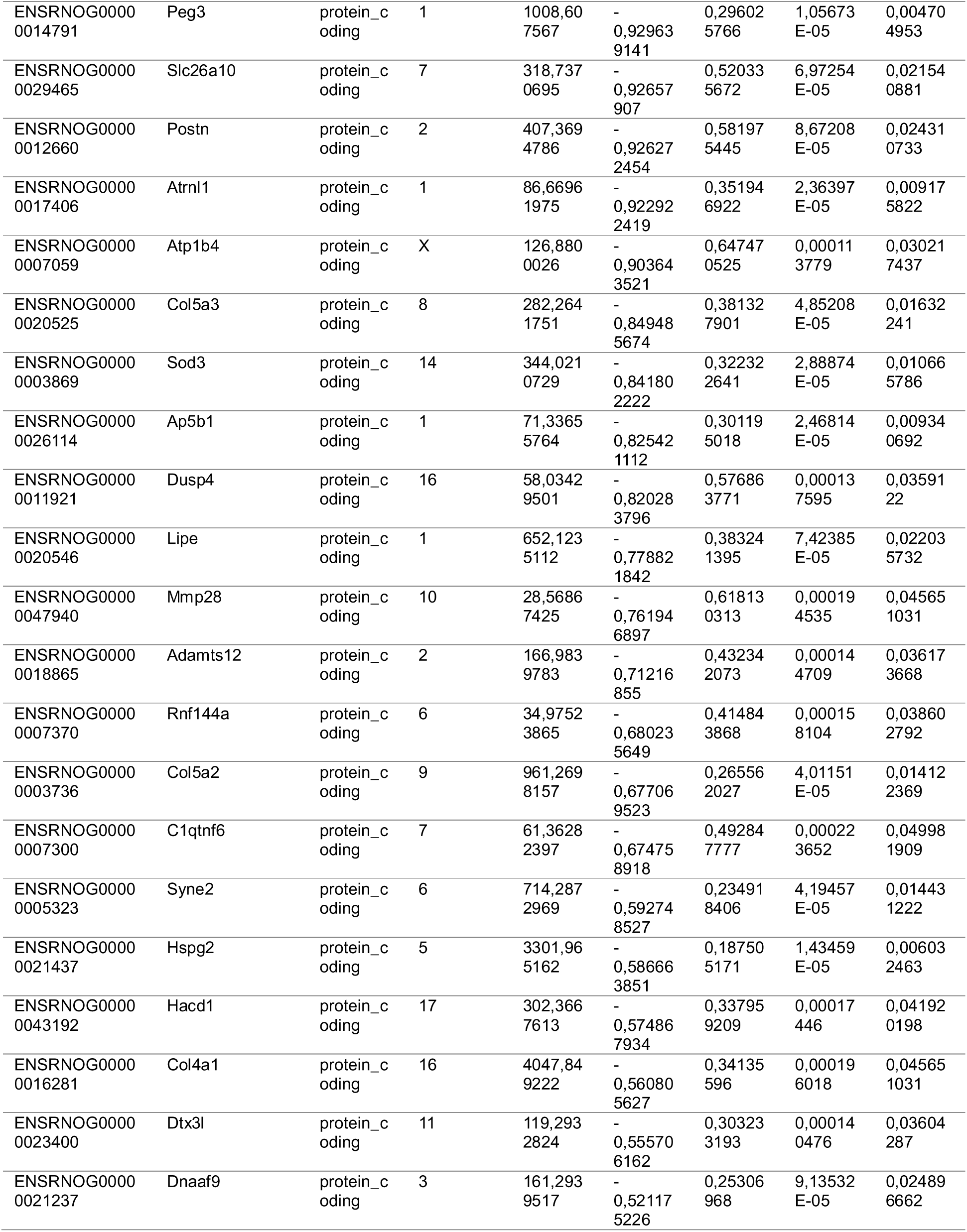
Differentially expressed genes (DEGs) altered by MyoMed-205 treatment compared with vehicle after 12 hours of UDD. From the 69 DEGs identified in DNV_205 vs. DNV_VEH listed above, 7 DEGs were upregulated (UP), while other 62 DEGs were downregulated (DOWN) in DNV_205 compared with DNV_VEH group. The DEGs are presented as UP and DOWN categories and ranked according to the highest to lowest fold change value. lfcSE, log fold change standard error), Padj, adjusted p-value.

Gene Set Enrichment Analysis (GSEA) was subsequently employed to identify biological pathways modulated by UDD and MyoMed-205, including hallmark pathways (Figure 1D), molecular functions (Figure 1E), and cellular compartments (Figure 1F). UDD significantly altered the expression of genes associated with muscle contraction, mitochondrial structure/function, inflammatory responses, proteolysis, apoptosis, and muscle tissue remodeling, compared with Sham. Interestingly, MyoMed-205 upregulated pathways involved with antioxidant response, chaperone-mediated unfolded protein response, ribosome biogenesis, protein synthesis, and cell growth. Conversely, MyoMed-205 downregulated genes associated with adipogenesis and extracellular matrix collagen deposition compared with DNV_VEH and Sham groups (Figure 1D-F).

### Effects of MyoMed-205 upon mechanisms that control skeletal muscle structure and contractile function

We previously reported that 12 hours of UDD promotes muscle fibers contractile dysfunction and atrophy [6]. Herein, our histomorphometric analysis confirmed a significant decrease in diaphragm fiber size across all types (Sham vs. DNV_VEH vs. DNV_205, mean ± SD - Type I: 1537 ± 269 vs. 1184 ± 91 vs. 1588 ± 342; Type IIa: 1750 ± 295 vs. 1307 ± 139 vs. 1795 ± 497; Type IIb/x: 3601 ± 788 vs. 2449 ± 308 vs. 3741 ± 1153 μm^2^, p < 0.05, *n* = 8) following 12 h of UDD (type I: p = 0.03; type IIa: p = 0.04; and type IIb/x: p = 0.02) compared with Sham. UDD caused comparable atrophy levels among the fiber types, despite a trend indicating a noticeable effect in fast-twitch type IIb/x fibers compared with slow-twitch type I fibers (p = 0.06). Fiber type relative distribution remained unchanged (Figure 2A-B). MyoMed-205 significantly protected against UDD-associated fiber atrophy (p < 0.05, *n* = 8), without showing fiber type-specific effect or affecting relative distribution (Figure 2A-B).

**Figure 2.**
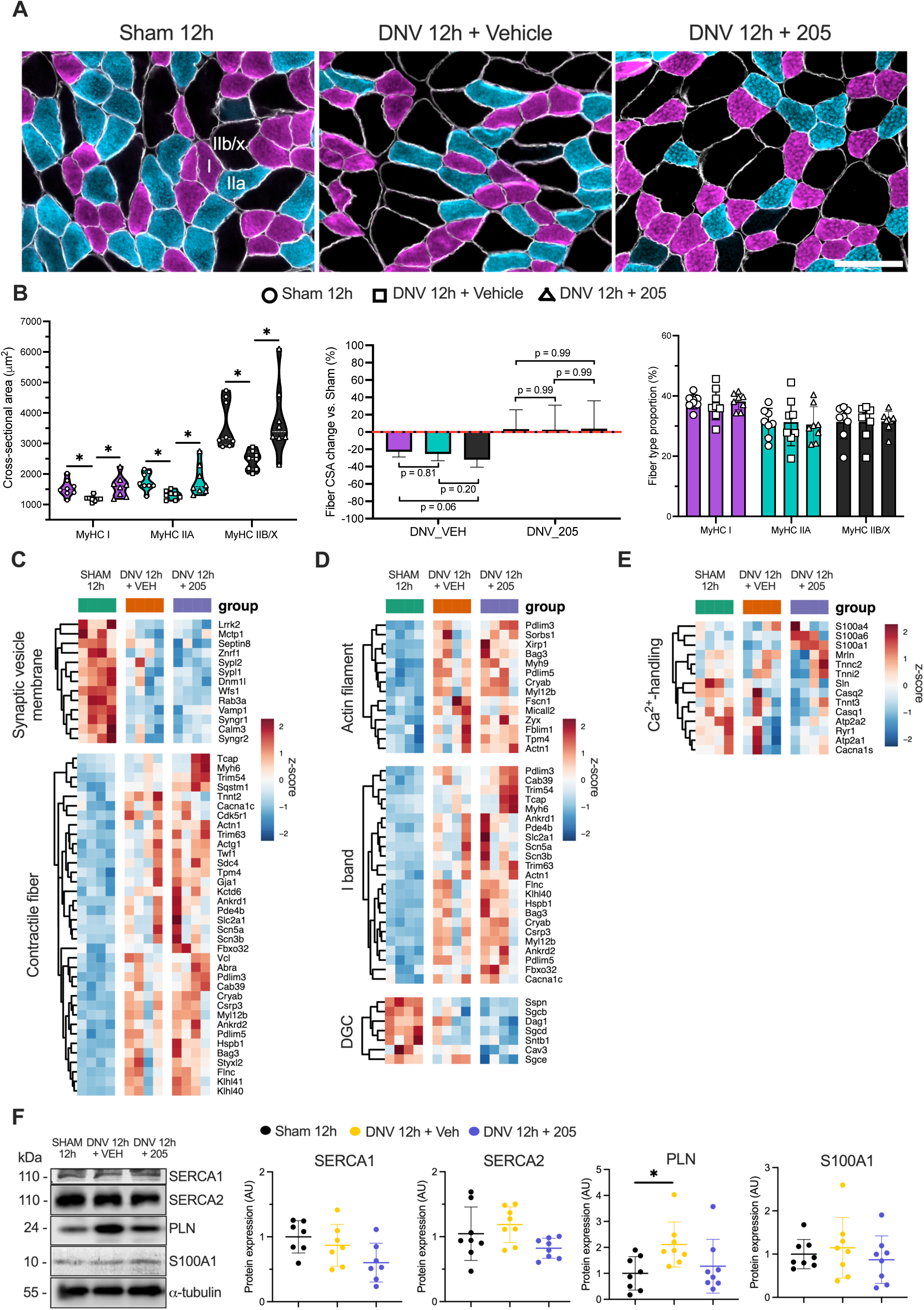
MyoMed-205 protects against diaphragm fiber atrophy induced by 12 hours of unilateral diaphragm denervation (UDD). (A) Representative photomicrographs of diaphragm muscle immunolabeled with antibodies against the myosin heavy chain (MyHC) isoforms, type I (violet), type IIa (light blue), type IIb/x (black), and dystrophin (white) for morphometric analysis by fiber type. Scale bar = 100 μm. (B) Fibers’ cross- sectional area (CSA), relative CSA percentual change compared with Sham-controls, and relative percentual distribution by fiber type. Data are presented as mean and standard deviation (*n* = 8). One-way ANOVA followed by Tukey’s post-hoc test was used for statistical comparisons among the groups. *, p < 0.05. (C) Heat map of genes linked to muscle contraction (i.e., synaptic vesicle membrane and contractile fiber). (D) Heat map of genes linked to sarcomere structure and function (i.e., actin filament, I band, and dystrophin glycoprotein complex (DCG)). (F) Protein levels of calcium- handling markers were assessed by Western blotting. Data are presented as mean and standard deviation of the fold change relative to the control group (*n* = 7 to 8). One-way ANOVA followed by Tukey’s post-hoc test was used for statistical comparisons among the groups. *, p < 0.05.

To gain mechanistic insights into the protective effects of MyoMed-205, we examined the expression levels of genes and proteins involved in the control of muscle structure and contraction. Significant changes in genes regulating synaptic vesicle membrane and muscle fiber structure/function were observed following 12 h of UDD (Figure 2C). Noteworthy, MyoMed-205 selectively enhanced the expression of genes involved with sarcomere organization and integrity, Z-disk structure, mechanotransduction, and intercellular communication (e.g., Tcap, Actn1, Pdlim3, Cryab, Ankrd1, Zyx, Sdc4, Sorbs1, Tpm4, Bag3 and Gja1) (Figure 2C-D).

Furthermore, UDD affected calcium handling pathways as indicated by downregulation of key calcium handling genes (e.g., RyR1, SERCA1, and SERCA2) (Figure 2E) and increased phospholamban protein levels compared with Sham (Figure 2F). MyoMed-205 enhanced S100A1 gene expression, a calcium-binding protein that positively regulates the activity of calcium channels, concomitantly with suppression of denervation-induced phospholamban protein expression (Figure 2E-F).

### UDD negatively affects the transcription of genes associated with mitochondrial dysfunction and oxidative stress, while MyoMed-205 enhances the transcription of antioxidant factors

Mitochondrial structure and function are essential for the maintenance of skeletal muscle function and homeostasis. Therefore, we assessed mitochondrial alterations following 12 h of UDD and MyoMed-205 in the diaphragm. Following 12 hours of UDD, significant changes were observed in the transcription of genes that control mitochondrial structure and function. Key mitochondrial biogenesis regulators (e.g., Ppargc1a) and dynamics mediators (e.g., Opa1, Mfn1, Mfn2) were markedly downregulated (Figure 3A). Genes encoding mitochondrial structural complexes (Figure 3B-C), as well as those regulating energy metabolism via oxidative phosphorylation and fatty acid oxidation (Figure 3D), were also markedly downregulated, further suggesting rapid changes in the expression of genes that regulate mitochondrial dynamics and function. Moreover, we observed increased transcriptional activation of the reactive oxygen species (ROS) pathway (Figure 3E). Interestingly, MyoMed-205 upregulated the expression of a subset of genes involved with the redox balance and antioxidant response (e.g., Nrf1, Nfe2l2, Sirt1, Sod2, Prdx1, Prdx6, and Gclc) (Figure 3E).

**Figure 3.**
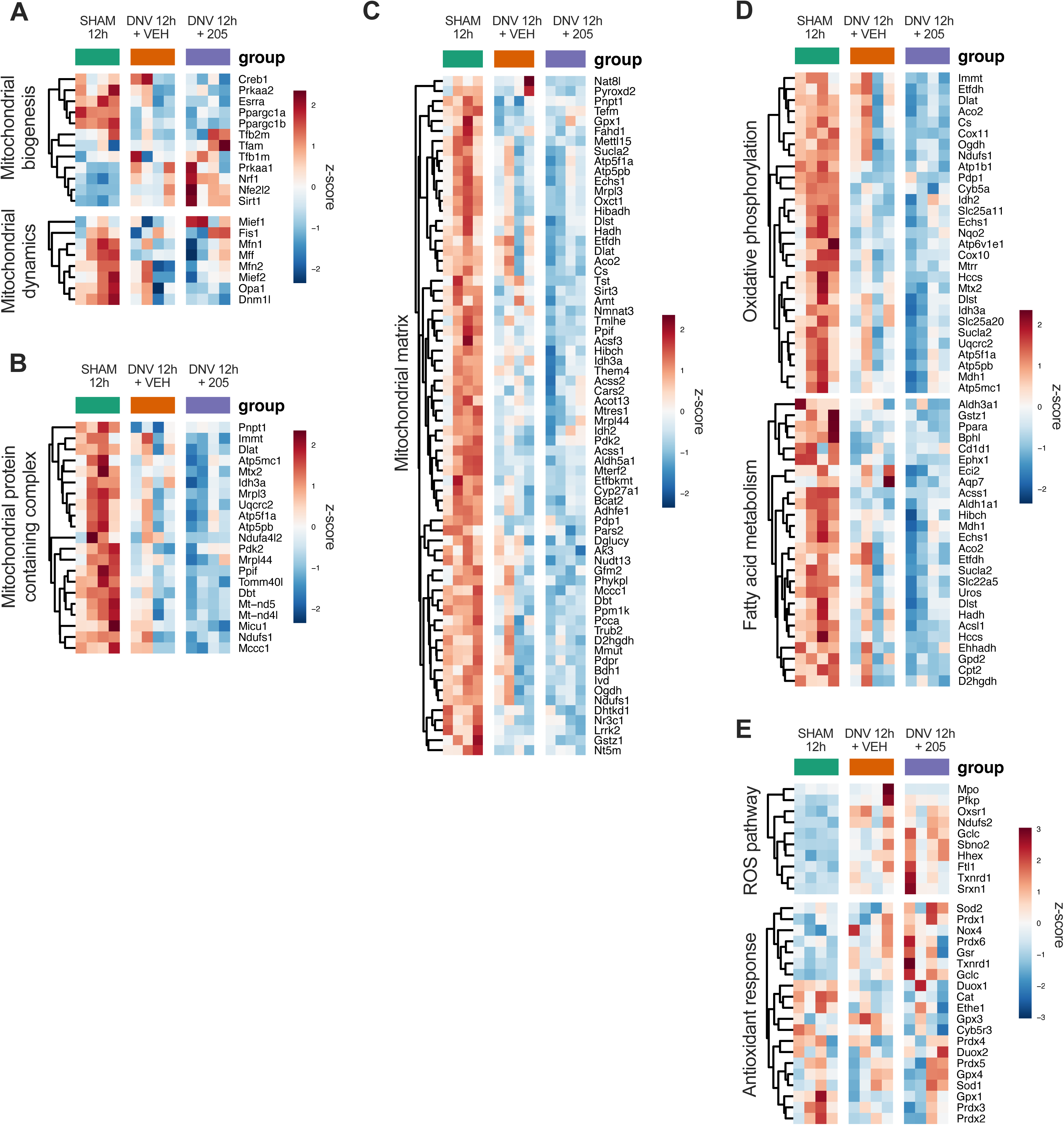
Unilateral diaphragm denervation (UDD) enhances transcriptional activation of pathways associated with mitochondrial dysfunction and oxidative stress, while MyoMed-205 upregulates the gene expression of antioxidant factors. (A) Heat map of genes involved in the control of mitochondrial biogenesis and dynamics. (B-C) Heat map of genes associated with mitochondrial structural and functional complexes. (D) Heat map of genes linked to mitochondrial energy production and metabolism. (E) Heat map of genes involved with reactive oxygen species (ROS) production and antioxidant response.

### MyoMed-205 enhances activation of pathways regulating ribosome biogenesis, protein synthesis, and cell growth under UDD-induced stress

Denervation can negatively affect muscle mass by altering protein synthesis rate and promoting negative protein net balance. We evaluated the impacts of UDD and the MyoMed-205 upon pathways involved with cell growth, proliferation, and survival. Noteworthy, Myc target genes (Figure 4A), the KRAS (Figure S3), and mTORC1 signaling pathways were found upregulated after 12 hours of UDD (Figure 4B). Strikingly, MyoMed-205 enhanced RNA processing and transcription (Figure S2), further amplifying these effects by increasing the expression of genes associated with the PI3K-Akt-mTOR pathway (Figure 4B), ribosome biogenesis, translation initiation (Figure 4C), and androgen response (Figure S3). Consistent with gene expression changes, MyoMed-205 increased PI3K-Akt-mTOR pathway activation at the protein level (Figure 4D).

**Figure 4.**
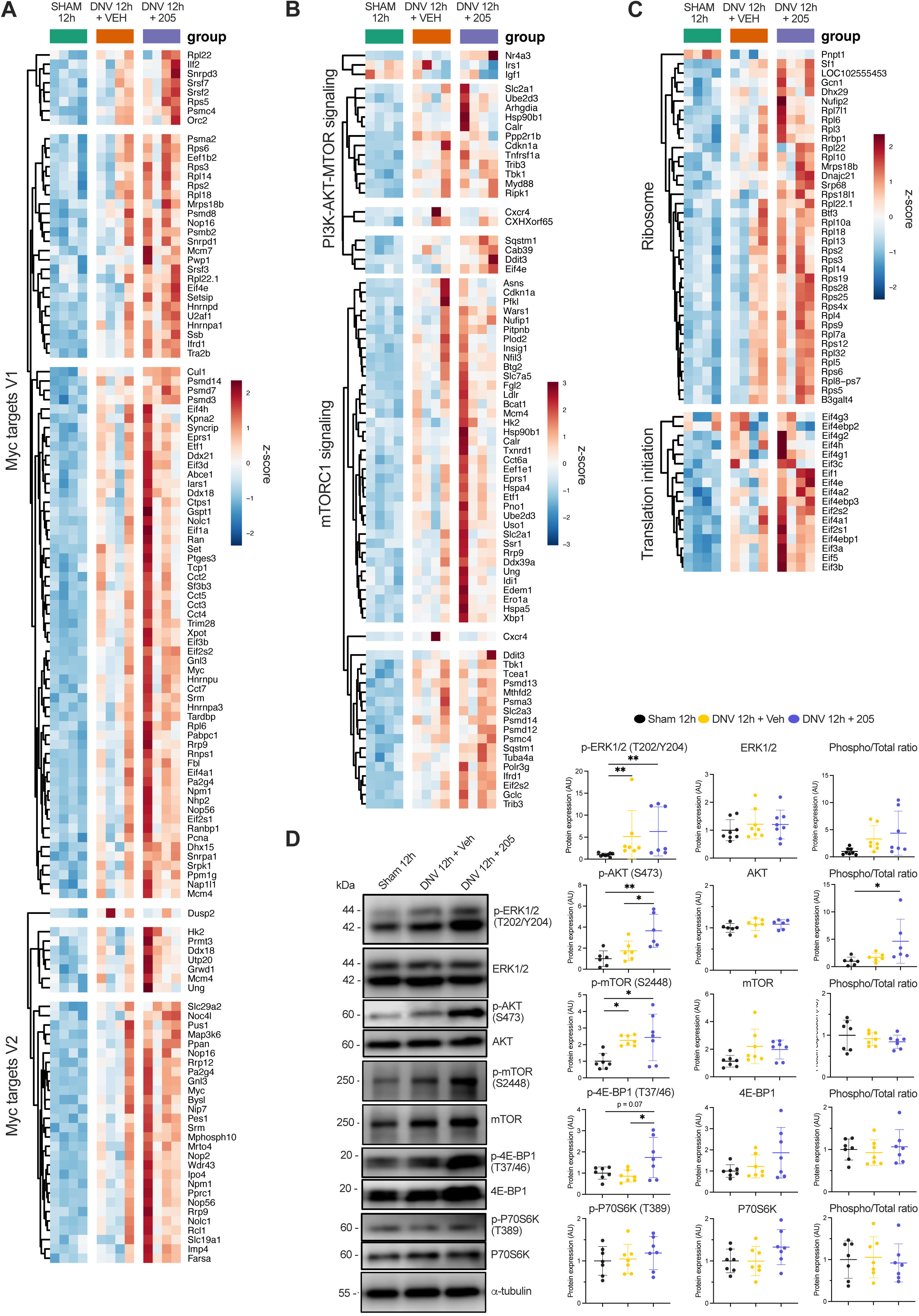
MyoMed-205 enhances the activation of signaling pathways controlling ribosome biogenesis, protein synthesis, and cell growth. (A) Heat map of Myc target genes. (B) Heat map of genes associated with the PI3K-AKT-mTOR pathway. (C) Heat map of genes controlling ribosome biogenesis and translation initiation. (D) Protein levels of MAPK and PI3K-Akt-mTOR pathways markers were assessed by Western blotting. Data are presented as mean and standard deviation of the fold change relative to the control group (*n* = 6 to 7). One-way ANOVA followed by Tukey’s post-hoc test was used for statistical comparisons among the groups. *, p < 0.05; **, p < 0.01.

### UDD upregulates proteolytic signaling pathways, while MyoMed-205 improves chaperone-mediated unfolded protein response at the transcriptional level

Denervation can also activate proteolytic pathways contributing to muscle wasting. Thus, we evaluated the effect of UDD and MyoMed-205 upon pathways associated with muscle protein breakdown and programmed cell death. UDD significantly upregulated distinct proteolytic and stress response pathways, including immune and inflammatory responses (Figure S4), the IL6-JAK-STAT3 pathway (Figure 5A), unfolded protein response (Figure 5B), ubiquitin-proteasome pathway (Figure 5C-D), p53 pathway (Figure 5E), and apoptosis (Figure 5F) at the transcriptional level. Remarkably, MyoMed-205 enhanced the expression of genes mediating chaperone-mediated unfolded protein response (Figure 5B). Noteworthy, we noticed a relatively higher gene expression of E3-ligases linked to muscle wasting, including Fbxo32 (i.e., Atrogin-1, log2fold = 0.96 vs 0.20, values relative to Sham; padj = 0.59) and TRIM63 (i.e., MuRF1, log2fold = 2.27 vs 1.44, values relative to Sham; padj = 0.59) in MyoMed-205 treated animals compared with vehicle, despite not reaching statistical significance (Figure 5C). In line with this, protein levels of these E3-ligases and K48-ubiquitinated proteins were not significantly changed, except for MuRF2, a close relative of MuRF1, which was upregulated by UDD but prevented by MyoMed-205 (Figure 5D).

**Figure 5.**
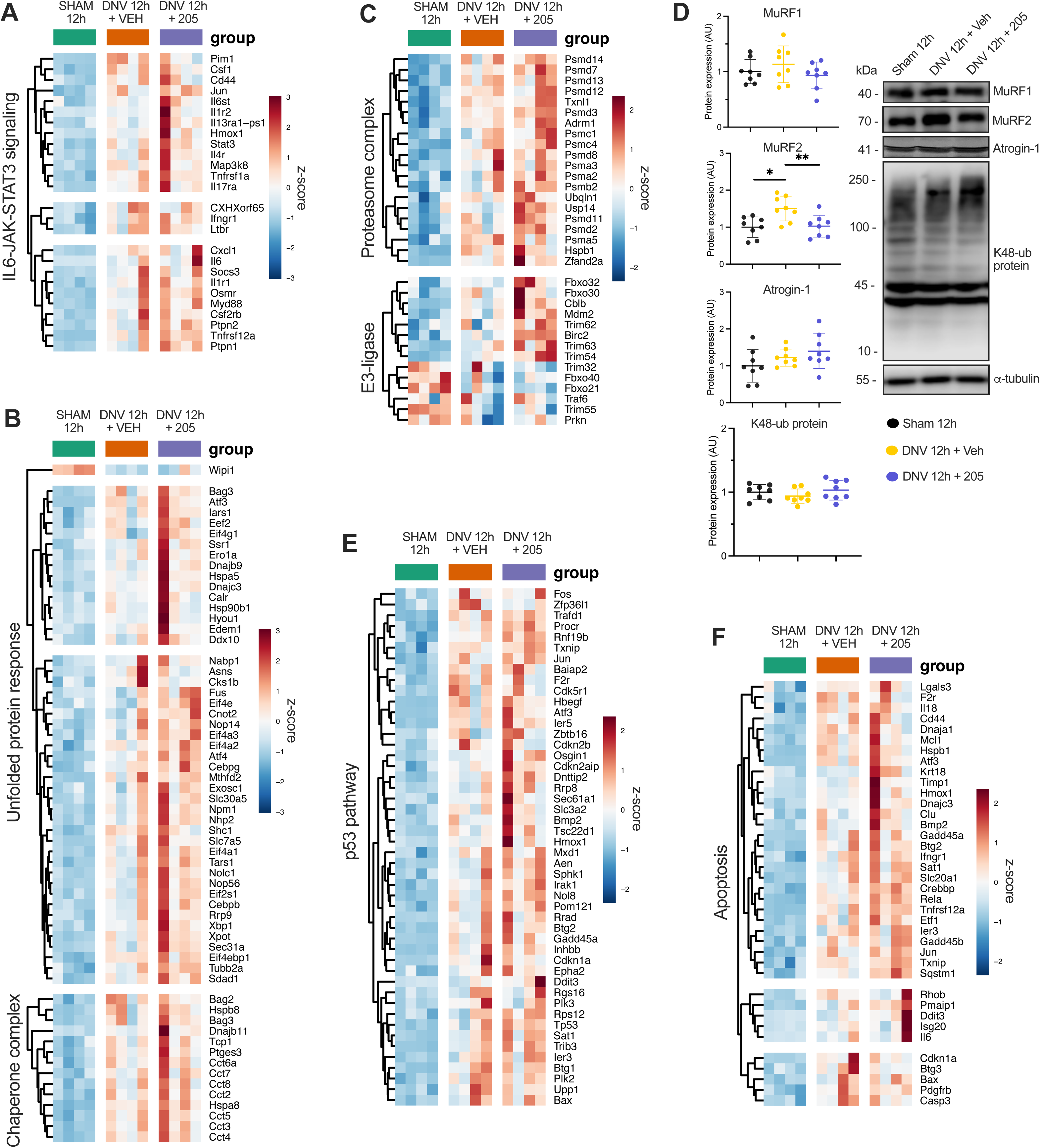
MyoMed-205 enhances the transcriptional activation of heat shock chaperones and the unfolded protein response (UPR) mechanism in the diaphragm under denervation-induced cellular stress and catabolic state. (A) Heat map of genes from the IL6-JAK-STAT3 pathway. (B) Heat map of genes associated with the chaperone complex and UPR. (C) Heat map of genes linked to the ubiquitin-proteasome system. (D) Protein levels of ubiquitin-proteasome pathway markers were assessed by Western blotting. Data are presented as mean and standard deviation of the fold change relative to the control group (*n* = 8). One-way ANOVA followed by Tukey’s post-hoc test was used for statistical comparisons among the groups. *, p < 0.05; **, p < 0.01. (E) Heat maps of genes from the p53 pathway. (F) Heat maps of genes associated with apoptosis.

### MyoMed-205 mitigates denervation-induced expression of genes involved with intramuscular lipid accumulation and ECM-collagen deposition

Prolonged inactivity can lead to sustained muscle cell stress and damage, triggering apoptosis and tissue remodeling responses such as intramuscular fat deposition and fibrosis. Therefore, we evaluated how UDD and MyoMed-205 affect these responses in the diaphragm after 12 hours.

Following UDD, we observed a significant upregulation in adipogenic-related gene expression compared with Sham (Figure 6A), although no changes were detected in adipogenic or lipid droplet-related proteins within this period (Figure 6B). Oil Red O (ORO) staining revealed significantly increased ORO-positive (ORO+) cells and intracellular lipid content across distinct fiber phenotypes (i.e., predominantly oxidative fibers <2500 μm², and predominantly glycolytic fibers >2500 μm²). Interestingly, MyoMed-205 downregulated adipogenesis at the transcriptional level (Figure 6A), attenuating intramuscular lipid accumulation triggered by UDD (Figure 6C).

**Figure 6.**
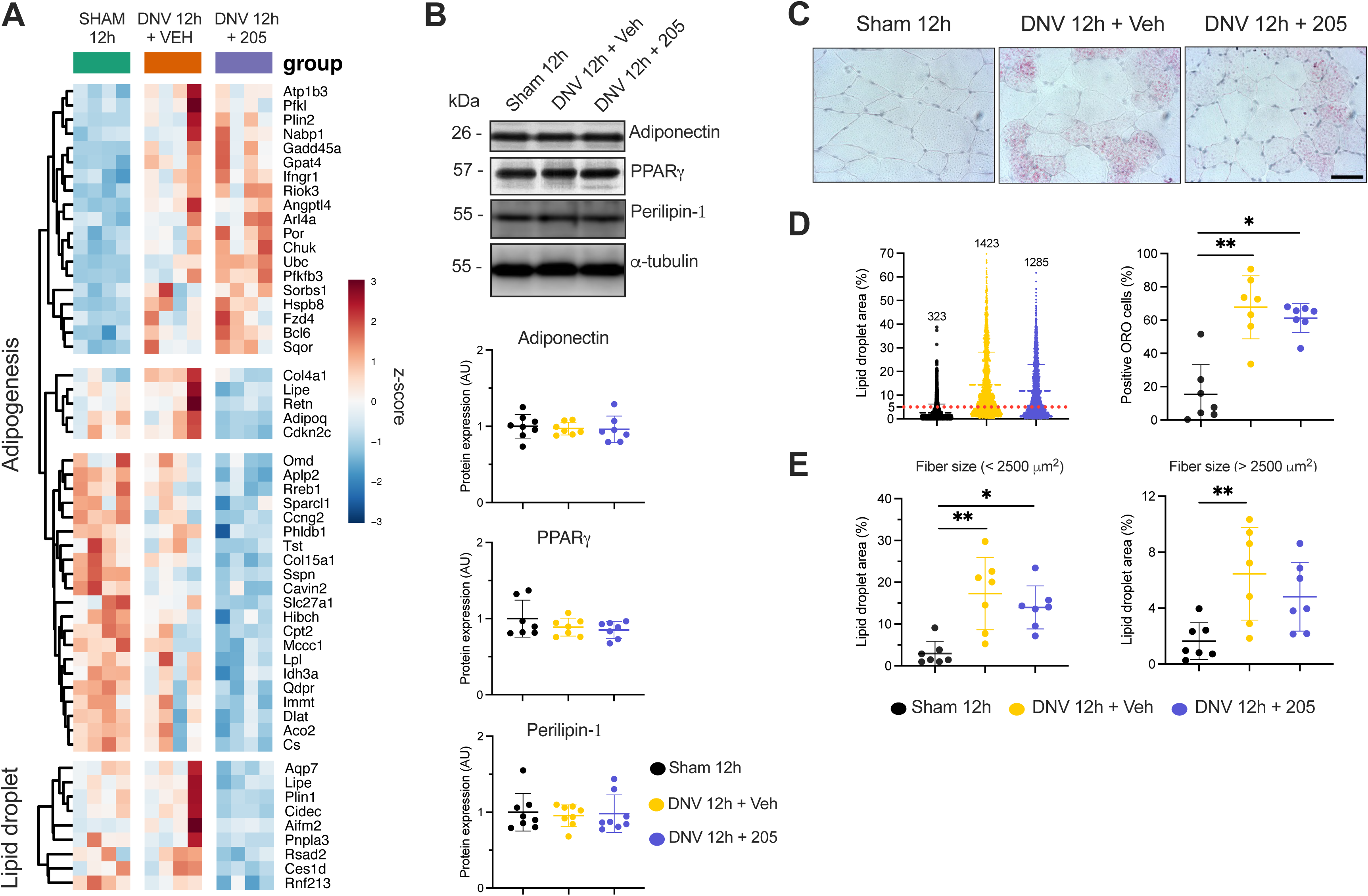
MyoMed-205 attenuates denervation-induced intramuscular fat accumulation in the diaphragm by downregulating pro-adipogenic genes. (A) Heat map of genes related to adipogenesis and intramuscular lipid accumulation. (B) Protein levels of adipogenesis pathway markers were assessed by Western blotting. Data are presented as mean and standard deviation of the fold change relative to the control group (*n* = 8). One-way ANOVA followed by Tukey’s post-hoc test was used for statistical comparisons among the groups. (C) Representative photomicrographs of Oil Red O (ORO) staining of the diaphragm for assessing intramuscular lipid accumulation following 12 h of unilateral diaphragm denervation and the effects of MyoMed-205. Scale bar = 50 μm. (D) Quantitative analysis of positive ORO muscle cells (i.e., intracellular lipid droplet area > 5% of total muscle fiber area). (E) Mean intracellular lipid droplet area of diaphragm muscle fibers < 2500 μm^2^ (predominantly oxidative metabolism) and fibers > 2500 μm^2^ (predominantly glycolytic metabolism). In total, 300 muscle cells were assessed per sample to estimate the average positive ORO cells and intracellular lipid content. Data are presented as mean and standard deviation (*n* = 7). One-way ANOVA followed by Tukey’s post-hoc test was used for statistical comparisons among the groups. *, p < 0.05; **, p < 0.01.

Moreover, UDD upregulated the expression of genes associated with the TGF-β signaling (Figure S5), collagen trimmer (Figure 7A), basement membrane (Figure 7B), and extracellular-matrix collagen deposition (Figure 7C), compared with Sham. On the other hand, MyoMed-205 downregulated the pro-fibrotic response at the transcriptional level (Figure 7A-C).

**Figure 7.**
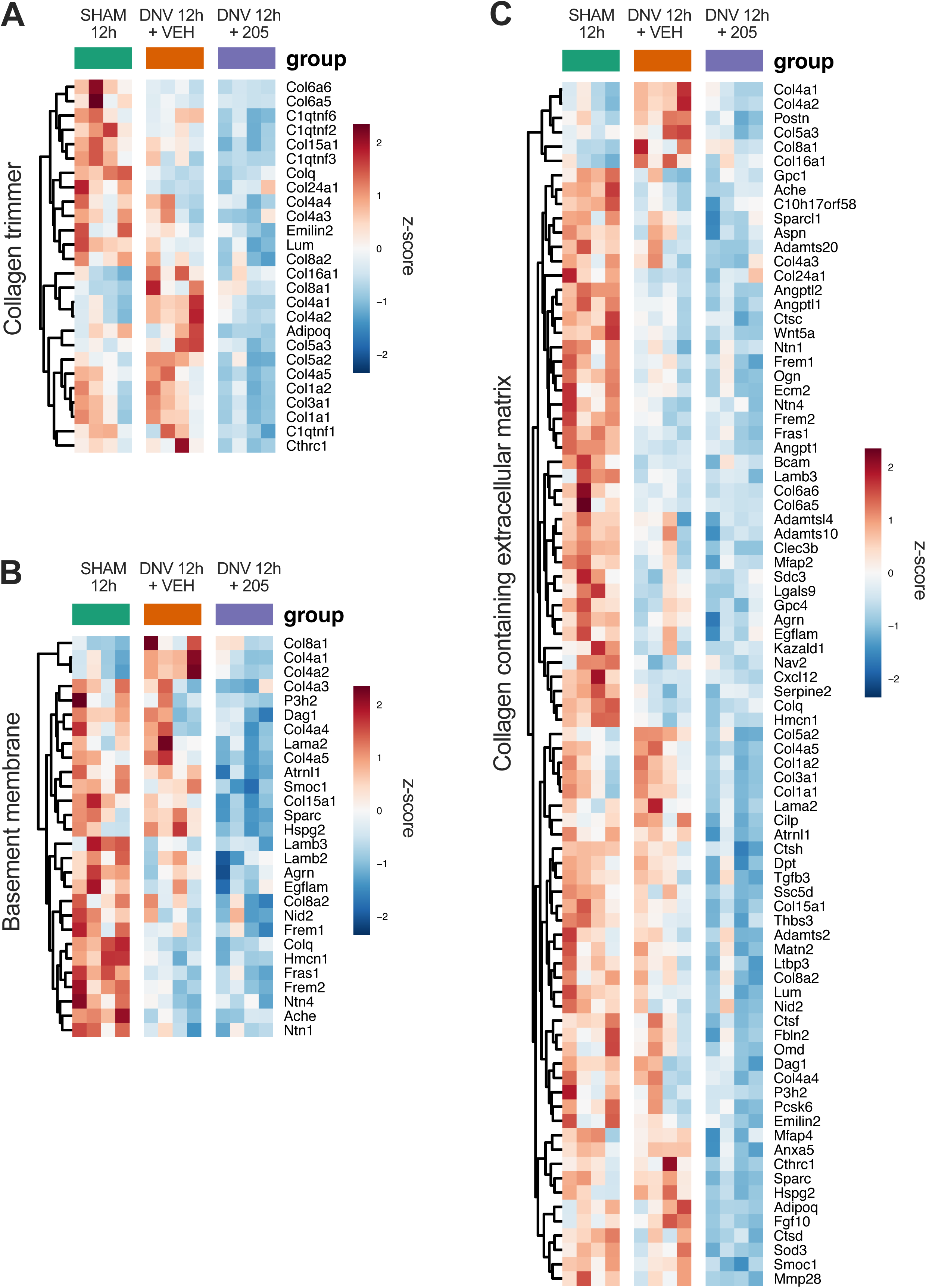
MyoMed-205 mitigates denervation-induced upregulation of genes involved with extracellular matrix collagen deposition and fibrosis in the diaphragm at the transcriptional level. (A) Heat map of genes involved in the formation of collagen trimmers. (B) Heat map of genes associated with the basement membrane. (C) Heat map of genes associated with collagen assembly and the extracellular matrix.

## Discussion

Lack of contractile activity can rapidly induce diaphragm muscle weakness and wasting, contributing to impaired respiratory function, morbidity, and mortality. This clinical condition is particularly found in patients undergoing phrenic nerve injury or prolonged mechanical ventilation during intensive care unit (ICU) treatment. Therefore, understanding the mechanisms underlying UDD and how to prevent them by identifying potential novel therapeutic targets is an important, and yet unmet, clinical need. We recently reported that small-molecule targeting MuRF1 (MyoMed-205) protects against early diaphragm muscle fiber contractile dysfunction and atrophy in rats undergoing 12 hours of UDD [6]. In the present study, we identified molecular mechanisms that are linked to the MyoMed-205-mediated protection against this loss in fibers’ contractile force and size. Briefly, MyoMed-205 upregulated pathways associated with: 1) muscle fiber structure and contractile function; 2) antioxidant defense; 3) chaperone-mediated unfolded protein response; 4) muscle cell growth, while downregulated pathways involved with 5) intramuscular lipid accumulation and 6) ECM collagen deposition at the transcriptional and/or protein level.

Disrupted calcium homeostasis has been reported in response diaphragm mechanical inactivity, as evidenced by downregulation of key genes like SERCA1, impaired RyR1 activity, decreased Ca2+ sensitivity, and sarcoplasmic reticulum (SR) calcium leakage [27]. Consistent with these findings, we observed a rapid downregulation of key genes involved in calcium handling, alongside increased phospholamban protein levels, a negative regulator of SERCA activity. MuRF1 has been reported to directly target proteins involved in calcium handling and muscle contraction [16–18]. Interestingly, MyoMed-205 prevented denervation-induced elevation of phospholamban protein levels, while upregulating S100A1 gene expression, although its protein levels remained unchanged. S100A1 is a calcium-binding protein that enhances RyR1 and SERCA activity and thus improves excitation-contraction coupling in striated muscles [28]. S100A1 overexpression has been shown to rescue cardiac function throughout enhanced Ca²⁺ handling and myofibrillar protein Ca²⁺ responsiveness in heart failure models [29]. These findings suggest that MyoMed-205 may partially preserve calcium handling homeostasis via upregulation of S100A1 and blockade of phospholamban, likely contributing to preserving diaphragm contractile function, as we previously reported [6].

Myofibrillar protein loss and post-translational modifications (PTMs) induced by prolonged inactivity can also impair diaphragm force-generation capacity. ROS-induced oxidative modifications of muscle proteins have been detected as early as 12 hours of UDD. Our data revealed rapid downregulation of key genes for mitochondrial function and augmented activation of oxidative stress pathways, consistent with our previous findings [6]. MuRF1 can directly regulate certain mitochondrial proteins [16–18]. Here, MyoMed-205 increased the mRNA levels of key antioxidant transcription factors (e.g., Nrf1, Nfe2l2) and enzymes (e.g., Sod2, Prdx1, Prdx6, and Gclc). These findings suggest that MyoMed-205 may improve antioxidant defense mechanisms, which likely contribute to protecting diaphragm muscle fiber structure and function under denervation stress.

Stress-induced unfolded protein response (UPR) activation contributes to maintaining muscle homeostasis under various stress states. Elevated oxidative stress and proteolysis contribute to myofibrillar protein damage, loss, and muscle wasting [30]. Ultrastructural changes, including impaired sarcomere and Z-disk structures, were observed in the diaphragm following denervation [8]. Accordingly, we observed increased UPR pathway activation, which indicates a cellular response against myofibrillar protein damage and loss after 12 hours of UDD. Interestingly, MyoMed-205 significantly enhanced UPR pathway activation at the transcriptional level by upregulating transcription factors (e.g., XBP1, ATF4) that regulate co-chaperone (e.g., Bag3) and chaperone complexes (e.g., Cryab, Hspa5, Hspa8, Hspb8, Hsp90b1, Dnjb11, Cct5, etc.). MyoMed-205 also upregulated the expression of genes involved in sarcomere integrity maintenance. Bag3 is known to facilitate chaperone-assisted selective autophagy (CASA) process for myofibrillar protein maintenance, and its downregulation impairs contractile function and protein turnover [31]. Bag3 can also interact with alpha B-crystallin (i.e., Cryab) to regulate I-band and Z-disk integrity, while its regulation of Hsp70 chaperone activity protects against damaged protein aggregates and muscle homeostasis [32]. Similarly, chaperone co-inducer BGP-15 attenuates ventilation-induced diaphragmatic dysfunction by activating Hsp72 and reducing myosin post-translational modifications [33]. These findings indicate that MyoMed-205-mediated protective effects might involve enhanced transcriptional activation of chaperones and UPR during UDD stress.

Muscle mass regulation crucially depends on the balance between protein synthesis and degradation. After 12 h of UDD, we observed comparable levels of atrophy across slow- and fast-twitch fibers, despite a trend toward a slightly greater atrophy in type IIb/x fibers was noticed, consistent with previous findings in similar durations (i.e., 24 hours) of diaphragm paralysis [7]. Importantly, some studies also reported no changes in slow- or fast-twitch fiber size at 24 hours of UDD [5]. On the other hand, fiber-type-specific responses seem more pronounced in long-term UDD. Rodents subjected to several days (e.g., 3 – 14) of UDD exhibit predominant atrophy of fast-twitch fibers, particularly of type IIb/x, whereas slow-twitch fibers, which are essential for basal respiratory function, appear less affected, unaltered, or display transient hypertrophy [5,8,9,12,34]. Yet, the long-term effects of UDD on diaphragm muscle fibers are intricate and remain not completely understood. Clinical observations illustrate this complexity as, in humans, prolonged UDD has been shown to induce progressive diaphragm fiber atrophy, with slow-twitch fibers appearing to be more rapidly and severely affected than fast-twitch fibers [35].

The decline in diaphragm fiber size after 12 hours of UDD indicates a rapid shift towards a predominantly catabolic state. Yet, we simultaneously observed an increased expression of genes linked to protein synthesis and muscle growth (e.g., Myc, MAPK/ERK, and mTORC1) following UDD, suggesting the activation of compensatory mechanisms to counteract muscle catabolism, in line with previous reports [10,11]. Importantly, MyoMed-205 prevented denervation-induced diaphragm fiber atrophy and enhanced gene expression of pathways associated with RNA processing, ribosome biogenesis, protein synthesis, and cell growth (e.g., Myc, Nr4a3, PI3K-Akt-mTOR, eIF4e), which is consistent with the evidences that MuRF1 can directly target ribosomal components and translation factors [16–18].

We previously have shown that MyoMed-205 increases Akt activation in the diaphragm under early UDD stress [6]. Herein, we further validated the MyoMed-205- mediated activation of the Akt-mTOR pathway at the protein level within 12 hours of UDD. Previous studies demonstrated that a murine MuRF1 gene knockout enhances Akt activation [26] and *de novo* protein synthesis. Thereby, MuRF1 KO mice seem less susceptible to muscle wasting [36]. Furthermore, MuRF1 can indirectly influence Myc activity by targeting Bin1, a negative regulator of Myc [18], which was found significantly upregulated in MyoMed-205 treated rats. MyoMed-205 also enhanced NR4A3 gene expression, which positively regulates mTORC1 signaling and contributes to exercise-induced muscle hypertrophy [37]. Our data suggests that MyoMed-205 activates anabolic pathways at the gene and protein levels that might contribute to counteract UDD-associated diaphragm muscle atrophy.

Increased protein damage and breakdown also promote muscle wasting. In this study, we found a significant upregulation of inflammatory and proteolytic pathways (e.g., IL6-JAK-STAT3, TNFα/NFκB, ubiquitin-proteasome system) in the diaphragm after 12 hours of UDD. Noteworthy, animals treated with MyoMed-205 showed relatively higher gene expression of MuRF1 (i.e., TRIM63) compared with DNV_VEH, although not statistically significant, and without affecting protein levels. This may suggest a compensatory cellular mechanism in an attempt to enhance MuRF1 expression and/or activity following denervation, consistent with previous studies [38]. This finding underscores the importance of dose-response optimization, as excessive MyoMed- 205 levels may disrupt muscle homeostasis and function, as we previously showed [6]. Conversely, we observed increased MuRF2 protein levels following 12 hours of UDD, a response abrogated by MyoMed-205, in line with previous studies suggesting that MuRF1 deletion/inhibition may downregulate MuRF2 expression, and that MyoMed-205 may also affect MuRF2 [22]. This effect may compromise MuRF1- MuRF2 heterodimerization and its cooperative role in myofibrillar protein ubiquitination [6,23,39]. Therefore, MyoMed-205-mediated protection against diaphragm fibers contractile dysfunction and atrophy might involve the disruption of MuRF1-MuRF2 cooperative role in myofibrillar breakdown.

Sustained cellular stress triggers myofiber apoptosis and diaphragm tissue remodeling, including intramuscular fat accumulation and fibrosis. Accordingly, we found upregulated transcription of adipogenic genes concomitantly with a higher number of ORO+ muscle cells with increased intracellular lipid content after 12 h of UDD. MyoMed-205 alleviated these effects, likely through downregulation of key adipogenic genes (e.g., Adiponectin, Pparγ). This might involve MyoMed-205 mediated blockade of MuRF2, known to suppress Pparγ expression/activity [40]. Furthermore, UDD enhanced gene expression linked to ECM collagen deposition and fibrosis, consistent with previous studies [13]. Notably, MyoMed-205 suppressed the denervation-induced ECM collagen deposition response at the transcriptional level. Previous experimental studies demonstrated that small-molecules targeting MuRF1 mitigate cardiac fibrosis, improving contractile function during heart failure [24]. Our data indicate that MyoMed-205 may contribute to the structural preservation of the diaphragm by mitigating denervation-induced intramuscular lipid accumulation and ECM collagen deposition triggered by UDD.

### Study Limitations

This is the first study to provide mechanistic insights into how the small-molecule targeting MuRF1 (MyoMed-205) protects against early denervation-induced diaphragmatic dysfunction and atrophy in an experimental model. However, the following limitations must be considered: 1) These analyses were performed at a single time point (12 hours), limiting assessment of temporal dynamics in small-molecule targeting MuRF1 effects. 2) This study was limited to male rats treated with a single dosage (50 mg/kg bw), precluding assessment of potential sex-specific differences or dose-response relationships in response to denervation and MyoMed- 205 treatment. 3) Our mechanistic findings rely primarily on transcriptomic and protein content analyses, which do not account for important regulatory mechanisms such as post-transcriptional and post-translational modifications or changes in protein activity. 4) Cause-and-effect experiments were not feasible at this time to validate which mechanisms are crucial for small-molecule targeting MuRF1-mediated protection during early UDD. Future studies should employ multi-omics analyses, assess different dosage ranges in both sexes, and evaluate multiple time points to further comprehend the complex molecular mechanisms underlying small-molecule targeting MuRF1-mediated protection against denervation-induced diaphragm weakness and wasting.

### Conclusion

The small-molecule targeting MuRF1, MyoMed-205, protects against denervation-induced diaphragm muscle contractile dysfunction and atrophy in rats undergoing 12 hours of UDD [6]. In this study, our mechanistic findings reveal that MyoMed-205 enhances the transcriptional mechanisms involved with the control of sarcomere structure and function, antioxidant defense, chaperone-mediated unfolded protein response, and muscle cell growth. MyoMed-205 also attenuated intramuscular lipid accumulation and pro-fibrotic responses triggered by UDD. These findings indicate MuRF1 as a promising therapeutic target for counteracting denervation-induced diaphragm muscle contractile dysfunction and atrophy.

## Author’s Contributions

FR and ASM were involved in the study’s conceptualization. FR conducted animal experiments, sample collection, functional, morphometric, and molecular analysis. PRJ conducted bioinformatic analysis (RNA-sequencing). FR and ASM performed data management and analysis. FR and ASM prepared the original manuscript. ASM and SL were involved in acquiring funding and project supervision. All authors were involved in the critical revision of the manuscript. All authors read and approved the final manuscript.

## Acknowledgments

We wish to thank Kelly Patricia for her excellent laboratory technical assistance.

## Ethical Standards Declaration

All animal studies have been approved by the appropriate ethics committee (CEUA ICB USP protocols #8728030320 and #5143091020), and have therefore been performed following the ethical standards laid down in the 1964 Declaration of Helsinki and its later amendments.

## References

1. Fogarty MJ, Mantilla CB, Sieck GC. Breathing: Motor Control of Diaphragm Muscle. Physiology 2018;33:113–126.

2. McCool FD, Tzelepis GE. Dysfunction of the Diaphragm. N Engl J Med 2012;366:932–942.

3. Gransee HM, Mantilla CB, Sieck GC. Respiratory muscle plasticity. Compr Physiol 2012;2:1441–1462.

4. Sieck GC, Fogarty MJ. Diaphragm Muscle: A Pump That Can Not Fail. Physiol Rev 2025 doi:10.1152/physrev.00043.2024.

5. Shindoh C, Hida W, Kurosawa H, Ebihara S, Kikuchi Y, Takishima T et al. Effects of unilateral phrenic nerve denervation on diaphragm contractility in rat. Tohoku J Exp Med 1994;173:291–302.

6. Ribeiro F, Alves PKN, Bechara LRG, Ferreira JCB, Labeit S, Moriscot AS. Small-Molecule Inhibition of MuRF1 Prevents Early Disuse-Induced Diaphragmatic Dysfunction and Atrophy. Int J Mol Sci 2023;24:3637.

7. Gill LC, Ross HH, Lee KZ, Gonzalez-Rothi EJ, Dougherty BJ, Judge AR et al. Rapid diaphragm atrophy following cervical spinal cord hemisection. Respir Physiol Neurobiol 2014;192:66–73.

8. Gosselin LE, Brice G, Carlson B, Prakash YS, Sieck GC. Changes in satellite cell mitotic activity during acute period of unilateral diaphragm denervation. J Appl Physiol 1994;77:1128–1134.

9. Aravamudan B, Mantilla CB, Zhan WZ, Sieck GC. Denervation effects on myonuclear domain size of rat diaphragm fibers. J Appl Physiol 2006;100:1617–1622.

10. Argadine HM, Hellyer NJ, Mantilla CB, Zhan WZ, Sieck GC. The effect of denervation on protein synthesis and degradation in adult rat diaphragm muscle. J Appl Physiol 2009;107:438–444.

11. Argadine HM, Mantilla CB, Zhan WZ, Sieck GC. Intracellular signaling pathways regulating net protein balance following diaphragm muscle denervation. Am J Physiol - Cell Physiol 2011;300:318–327.

12. van der Pijl R, Strom J, Conijn S, Lindqvist J, Labeit S, Granzier H et al. Titin-based mechanosensing modulates muscle hypertrophy. J Cachexia Sarcopenia Muscle 2018;9:947–961.

13. Gosselin LE, Sieck GC, Aleff RA, Martinez DA, Vailas AC. Changes in diaphragm muscle collagen gene expression after acute unilateral denervation. J Appl Physiol 1995;79:1249–1254.

14. Centner T, Yano J, Kimura E, McElhinny AS, Pelin K, Witt CC et al. Identification of muscle specific ring finger proteins as potential regulators of the titin kinase domain. J Mol Biol 2001;306:717–726.

15. Bodine SC, Latres E, Baumhueter S, Lai VK-M, Clarke BA, Poueymirou WT et al. Identification of ubiquitin ligases required for skeletal muscle atrophy. Science (80-) 2001;294:1704–1708.

16. Witt SH, Granzier H, Witt CC, Labeit S. MURF-1 and MURF-2 target a specific subset of myofibrillar proteins redundantly: Towards understanding MURF-dependent muscle ubiquitination. J Mol Biol 2005;350:713–722.

17. Witt CC, Witt SH, Lerche S, Labeit D, Back W, Labeit S. Cooperative control of striated muscle mass and metabolism by MuRF1 and MuRF2. EMBO J 2008;27:350– 360.

18. Baehr LM, Hughes DC, Lynch SA, Van Haver D, Maia TM, Marshall AG et al. Identification of the MuRF1 Skeletal Muscle Ubiquitylome Through Quantitative Proteomics. Function 2021;2:1–18.

19. Peris-Moreno D, Taillandier D, Polge C. MuRF1/TRIM63, Master Regulator of Muscle Mass. Int J Mol Sci 2020;21:6663.

20. Hooijman PE, Beishuizen A, Witt CC, de Waard MC, Girbes ARJ, Spoelstra-de Man AME et al. Diaphragm muscle fiber weakness and ubiquitin–proteasome activation in critically ill patients. Am J Respir Crit Care Med 2015;191:1126–1138.

21. Liu J, Chen Y, Han D, Huang M. Inhibition of the expression of TRIM63 alleviates ventilator-induced diaphragmatic dysfunction by modulating the PPARα/PGC-1α pathway. Mitochondrion 2025;83:102025.

22. Bowen TS, Adams V, Werner S, Fischer T, Vinke P, Brogger MN et al. Small-molecule inhibition of MuRF1 attenuates skeletal muscle atrophy and dysfunction in cardiac cachexia. J Cachexia Sarcopenia Muscle 2017;8:939–953.

23. Adams V, Bowen TS, Werner S, Barthel P, Amberger C, Konzer A et al. Small-molecule-mediated chemical knock-down of MuRF1/MuRF2 and attenuation of diaphragm dysfunction in chronic heart failure. J Cachexia Sarcopenia Muscle 2019;10:1102–1115.

24. Adams V, Schauer A, Augstein A, Kirchhoff V, Draskowski R, Jannasch A et al. Targeting MuRF1 by small molecules in a HFpEF rat model improves myocardial diastolic function and skeletal muscle contractility. J Cachexia Sarcopenia Muscle 2022;13:1565–1581.

25. Adams V, Gußen V, Zozulya S, Cruz A, Moriscot A, Linke A et al. Small-Molecule Chemical Knockdown of MuRF1 in Melanoma Bearing Mice Attenuates Tumor Cachexia Associated Myopathy. Cells 2020;9:2272.

26. Labeit S, Hirner S, Bogomolovas J, Cruz A, Myrzabekova M, Moriscot A et al. Regulation of Glucose Metabolism by MuRF1 and Treatment of Myopathy in Diabetic Mice with Small Molecules Targeting MuRF1. Int J Mol Sci 2021;22:2225.

27. Matecki S, Dridi H, Jung B, Saint N, Reiken SR, Scheuermann V et al. Leaky ryanodine receptors contribute to diaphragmatic weakness during mechanical ventilation. Proc Natl Acad Sci U S A 2016;113:9069–9074.

28. Kiewitz R, Acklin C, Schäfer BW, Maco B, Uhrík B, Wuytack F et al. Ca2+-dependent interaction of S100A1 with the sarcoplasmic reticulum Ca2+-ATPase2a and phospholamban in the human heart. Biochem Biophys Res Commun 2003;306:550– 557.

29. Most P, Bernotat J, Ehlermann P, Pleger ST, Reppel M, Börries M et al. S100A1: A regulator of myocardial contractility. Proc Natl Acad Sci 2001;98:13889–13894.

30. Shanely RA, Zergeroglu MA, Lennon SL, Sugiura T, Yimlamai T, Enns D et al. Mechanical ventilation–induced diaphragmatic atrophy Is associated with oxidative injury and increased proteolytic activity. Am J Respir Crit Care Med 2002;166:1369– 1374.

31. Martin TG, Myers VD, Dubey P, Dubey S, Perez E, Moravec CS et al. Cardiomyocyte contractile impairment in heart failure results from reduced BAG3-mediated sarcomeric protein turnover. Nat Commun 2021;12:2942.

32. Meister-Broekema M, Freilich R, Jagadeesan C, Rauch JN, Bengoechea R, Motley WW et al. Myopathy associated BAG3 mutations lead to protein aggregation by stalling Hsp70 networks. Nat Commun 2018;9:5342.

33. Salah H, Li M, Cacciani N, Gastaldello S, Ogilvie H, Akkad H et al. The chaperone co-inducer BGP-15 alleviates ventilation-induced diaphragm dysfunction. Sci Transl Med 2016;8:350ra103.

34. Geiger PC, Cody MJ, Macken RL, Bayrd ME, Sieck GC. Effect of unilateral denervation on maximum specific force in rat diaphragm muscle fibers. J Appl Physiol 2001;90:1196–1204.

35. Welvaart WN, Paul MA, van Hees HWH, Stienen GJM, Niessen JWM, de Man FS et al. Diaphragm muscle fiber function and structure in humans with hemidiaphragm paralysis. Am J Physiol - Lung Cell Mol Physiol 2011;301:228–235.

36. Koyama S, Hata S, Witt CC, Ono Y, Lerche S, Ojima K et al. Muscle RING-Finger Protein-1 (MuRF1) as a connector of muscle energy metabolism and protein synthesis. J Mol Biol 2008;376:1224–1236.

37. Pillon NJ, Gabriel BM, Dollet L, Smith JAB, Sardón Puig L, Botella J et al. Transcriptomic profiling of skeletal muscle adaptations to exercise and inactivity. Nat Commun 2020;11:470.

38. Furlow JD, Watson ML, Waddell DS, Neff ES, Baehr LM, Ross AP et al. Altered gene expression patterns in muscle ring finger 1 null mice during denervation- and dexamethasone-induced muscle atrophy. Physiol Genomics 2013;45:1168–1185.

39. Nguyen T, Bowen TS, Augstein A, Schauer A, Gasch A, Linke A et al. Expression of MuRF1 or MuRF2 is essential for the induction of skeletal muscle atrophy and dysfunction in a murine pulmonary hypertension model. Skelet Muscle 2020;10:1–10.

40. He J, Quintana MT, Sullivan J, L Parry T, J Grevengoed T, Schisler JC et al. MuRF2 regulates PPARγ1 activity to protect against diabetic cardiomyopathy and enhance weight gain induced by a high fat diet. Cardiovasc Diabetol 2015;14:97.

