## Supplementary information for "Small-molecule targeting MuRF1 protects against denervation-induced diaphragmatic dysfunction: Underlying molecular mechanisms"

### Material and Methods

#### Animals

Wistar rats (Male, 2-3 months old) were used in this study. The animals were kept in standard cages under controlled environmental conditions (24 ± 1°C, 12 h/12 h light-dark cycle) with access to standard food and water ad libitum. This study was approved and followed the institutional guidelines for animal care and use for research (CEUA ICB USP #8728030320 and #5143091020).

#### Experimental design

Twenty-four rats were randomly allocated into three experimental groups ($n$ = 8): Sham 12h (Sham); unilateral diaphragm denervation 12 hours + vehicle (DNV + VEH); and unilateral diaphragm denervation 12 hours + MyoMed-205 (DNV + 205). After 12 hours, the animals were euthanized under anesthesia, and the denervated right diaphragm muscle (costal portion) was harvested, quickly snap-frozen in 2-methylbutane chilled in liquid nitrogen, and stored at -80°C until further analysis. Diaphragm samples used in this study originate from our previous work [1].

#### MyoMed-205: formulation and delivery

MuRF1 inhibitor (MyoMed-205, Myomedix GmbH, Germany) was solubilized in an 8 mL vehicle solution consisting of DMSO-PEG400-Saline 0.9% (20% - 50% - 30%, respective proportions). The formulations were freshly prepared immediately before the animal experiments. For this, the compound was first supplemented with the calculated volume of DMSO, vortexed for 1 min, and sonicated at ~40°C for 2 minutes. Next, the corresponding volumes of PEG400 and Saline 0.9% were added to the formulation, vortexed for 1 min, and sonicated at ~40°C for 2 minutes. The delivery of either vehicle (VEH) or MyoMed-205 (205) treatments was managed in four split doses (2 mL/dose) administered every 3 h interval over the 12-hour experimental period via the caudal vein. Treatment started immediately following the confirmation of unilateral diaphragm paralysis [1].

**Unilateral diaphragm denervation**

To induce unilateral diaphragm denervation, the rats were initially sedated and anesthetized via intraperitoneal administration of acepromazine (2.5 mg/kg bw), followed after 30 minutes by the administration of ketamine (100 mg/kg bw) and xylazine (5 mg/kg bw) solution [1]. After reaching the surgical plane, the right phrenic nerve was exposed and transected at the rat's lower neck region. The absence of right diaphragm dome contractile activity was used to confirm the denervation's efficacy. Then, the surgical wound was sutured and treated with a topical antiseptic solution. During the post-operative period, animals were allowed to recover from anesthesia in individual cages with *ad libitum* access to food chow and water.

**Diaphragm muscle fiber typing and size analysis**

Diaphragm muscle cryosections (8 μm thick) were prepared using a cryostat at −20°C (Leica CM1850, Germany). For histomorphometric analysis by fiber type, frozen cryosections were allowed to thermoequilibrate for 30 minutes at room temperature. Next, samples were washed with PBS solution (1 x 5 minutes), incubated with M.O.M. blocking reagent (MKB-2213-1, VectorLabs) for 1 hour at room temperature. Next, samples were washed with PBS (3 x 5 minutes), and then incubated with primary antibodies overnight at 4°C (Table S1). Next day, samples were washed with PBS (3 x 5 minutes), and the secondary antibodies were incubated for 1 h at 37°C (Table S1).

Next, samples were washed with PBS (3x 5 minutes), and the slides were finished with mounting medium. Photomicrographs were acquired using Axio Scope A1 microscope (Zeiss, Germany). Diaphragm fibers' type distribution and cross-sectional area (CSA) were assessed with ImageJ (Fiji, NIH, USA). Representative muscle fiber type distribution and CSA values were determined as the average from 300 measurements per sample of previously unanalyzed muscle fibers [1].

**Oil Red O Staining (ORO)**

Diaphragm muscle cryosections (8 µm thick) were initially fixed using 10% PFA solution for 10 minutes, then rinsed with dH20 and 60% isopropyl alcohol solutions. Next, tissues were stained with Oil Red O (Sigma #O0635) working solution for 30 minutes and rinsed with dH20 and 60% isopropyl alcohol solutions. Tissues were incubated with Mayer's hematoxylin solution for 10 minutes for cell nuclei staining. After rinsing with dH20, slides were mounted using glycerine jelly aqueous-based mounting medium and photomicrographs acquired using Axio Scope A1 microscope (Zeiss, Germany). Representative number of positive ORO (ORO+) muscle cells and intracellular lipid content were determined based on measurements of 300 fibers per sample.

**Transcriptomic profiling (RNA-seq)**

Total RNA from diaphragm was extracted using TRIzol (Life Technologies, Carlsbad, CA). RNA integrity was analyzed in a Bioanalyzer 2100 (Agilent, Santa Clara, CA, USA). Four samples of each experimental group with an average 260/280nm optic density ratio higher than 1,8 and the RNA integrity number (RIN) higher than 8.0 (i.e., Sham 2, Sham 4, Sham 5, Sham 7, DNV_VEH 3, VEH 4, VEH 6, VEH 8, DNV_205 5,

205 6, 205 7, and 205 8) were selected for RNA-seq and transcriptomic analysis (See Figure Sx). Approximately ~1 ug of sample total RNA was used for cDNA library preparation performed according to the Illumina mRNA stranded poly-A tail enrichment Kit manufacturer's instructions (Illumina, San Diego, CA). RNA sequencing was carried out using the Illumina NextSeq2000 (NextSeq 2x100pb, 20 million paired-end reads depth). The RNA-req procedures were performed at NGS Soluções Genômicas (Piracicaba, Brazil).

RNA-seq bioinformatic analysis was performed similarly to previous reports [2]. Quality control of raw reads was performed using the FastQC toolkit (Babraham Bioinformatics). The reads were then aligned to the mRatBN7.2 rat genome using STAR [3]. Feature count was performed using featureCounts [4], and differential expression analyses were generated in R using the DESeq2 package [5] with adaptive log-fold change shrinkage estimator from the ashr package [6]. Pathway analysis was carried out by Gene Set Enrichment Analysis (GSEA) using a pre-ranked list of genes by log2Fold-Change and Molecular Signatures Database [7,8] and hallmark pathways gene sets [9], for pathway analysis. Transcriptome-related raw files and data were deposited in the GEO repository (GSE305304).

**Western Blotting**

Samples were powdered in a liquid nitrogen-chilled mortar and homogenized in RIPA buffer enriched with protease and phosphatase inhibitors (1 mM EDTA, pH 7.4, 0.625% sodium deoxycholate, 0.625% nonidet P-40, 6.2 mM sodium phosphate and protease and phosphatase inhibitor cocktail) (Thermo Fisher Scientific, Category Number #78445, Rockford, IL, USA). Then, homogenates were incubated on ice for

10 minutes, centrifuged at 10,000 × g for 10 minutes at 4°C. Next, supernatant containing the total protein was collected and stored at −80°C until further analysis.

Protein concentration was determined by the Bradford method with bovine serum albumin (BSA) concentration solution curve as the standard. Total protein was loaded onto an 8–15% polyacrylamide gel (SDS-PAGE) and separated via electrophoresis (constant 100 V). Then, proteins were transferred to a polyvinylidene difluoride membrane 0.45 µm (Thermo Fisher Scientific, #88518, Rockford, IL, USA) using a wet-transfer system (100 V, 350 mA, for 120 min). To verify a homogeneous loading and transfer efficacy, membranes were stained with Ponceau red solution. Next, membranes were washed with TBS-T 0.1% (0.5 M NaCl, 50 mM Tris-HCl pH 7.4, 0.1% Tween 20) (3 x 5 minutes), and blocked with 5% BSA or dry-fat milk in TBS-T 0.1% for 1 hour at room temperature. Subsequently, membranes were washed with TBS-T 0.1% (3 x of 5 minutes) and incubated with primary antibodies overnight at 4°C (see Table S1). Next, membranes were washed with TBS-T0.1% (3 rounds of 5 minutes), and incubated with the appropriate secondary antibody (Table S1). Membranes were stripped and re-probed for multiple protein detection within the same membrane. For this, membranes were incubated with mild stripping buffer (0.2 M glycine, 0.1% SDS (w/v), 1% Tween 20 (v/v), pH 2.2) (2 x 5 minutes), washed with PBS (2 x 10 minutes), and additionally with TBS-T0.1% (2 x 5 minutes), before returning to the blockage stage. Finally, membranes were washed with TBS-T 0.1% (3 x 5 minutes) and incubated with ECL substrate (Immobilon Forte, Millipore #WBLUF0500) for 5 minutes before signal detection acquired using C-digit blot scanner (Li-Cor, USA), or with BCIP/NBT substrate (Sigma #B1911) for colorimetric detection. Alpha-tubulin was used as a loading control for data normalization, and densitometry analysis was performed using ImageStudio software. Data are presented as fold-change relative to Sham control protein levels.

**Statistical Analysis**

Data analysis was performed Excel, Prism and R. For the analysis of the transcriptomic data, refer to the respective section above and supplementary information. For the analysis of histological, morphometric, and protein expression levels, the Shapiro-Wilk test was used to assess data normality. One-way ANOVA followed by Tukey's post-hoc test, or the Kruskal-Wallis followed by Dunn's test, were used for multiple comparisons between groups, for parametric and non-parametric data, respectively. Data are presented as mean ± standard deviation and statistical significance was accepted as $p < 0.05$.

**Supplementary Table Legends**

Table S1. List of antibodies used for immunofluorescence and western blotting analysis. IF, immunofluorescence; WB, western blotting. Antibodies were diluted in a 5% BSA PBS solution during the immunofluorescent assay. During western blotting assays, antibodies were diluted in a 5% BSA or non-fat dry milk TBS-T 0.1% solution.

Table S2. Quality control parameters of the RNA isolated from diaphragm muscles. Sham, Sham-operated 12 h controls; DNV_VEH, denervation 12 h + vehicle treatment; DNV_205, denervation 12 h + MyoMed-205 treatment; RIN, RNA integrity number; NA, not applicable.

**Supplementary Figure Legends**

Figure S1. Quality control analysis of the RNA samples isolated from the diaphragm for RNA-seq-based global gene expression profiling. (A) RNA ScreenTape quality control analysis and samples' RNA integrity number (RIN) scores. The four RNA samples of each experimental group with higher purity and integrity scores that were selected for RNA-seq transcriptomic analysis are highlighted in red. (B) RNA-seq quality control assessment of the number of genes detected, mitochondrial, AND ribosomal gene proportions. (C) RNA-seq sample distances analysis representing the overall gene expression similarities or differences among the samples.

Figure S2. MyoMed-205 upregulates RNA processing and transcription under 12 hours of unilateral diaphragm denervation. (A) Heat map of genes associated with the Cajal Body. (B) Heat map of genes associated with the nuclear speck. (C) Heat map of genes associated with the spliceosome complex. (D) Heat map of genes associated with the transcription export complex.

Figure S3. Effects of unilateral denervation and MyoMed-205 upon non-canonical signaling pathways involved with muscle cell growth and survival in the diaphragm. (A) Heat map of genes involved in KRAS signaling that were upregulated by denervation in the diaphragm. (B) Heat map of genes involved in KRAS signaling that were downregulated by denervation in the diaphragm. (C) Heat map of genes involved in androgen response that were upregulated by MyoMed-205.

Figure S4. Denervation enhances inflammatory and immune responses in the diaphragm. (A) Heat map of genes associated with TNF$\alpha$ signaling via NF$\kappa$B. (B) Heat map of genes associated with inflammatory response. (C) Heat map of genes associated with the complement system. (D) Heat map of genes associated with interferon gamma response.

Figure S5. Denervation enhances TGF-β signaling in the diaphragm. Heat map of genes associated with the TGF-β-signaling that were modulated following 12 h of unilateral diaphragm denervation.

### References


1. Ribeiro F, Alves PKN, Bechara LRG, Ferreira JCB, Labeit S, Moriscot AS. Small-Molecule Inhibition of MuRF1 Prevents Early Disuse-Induced Diaphragmatic Dysfunction and Atrophy. *Int J Mol Sci* 2023;**24**:3637.
2. Correia JC, Jannig PR, Gosztyla ML, Cervenka I, Ducommun S, Præstholm SM *et al.* Zfp697 is an RNA-binding protein that regulates skeletal muscle inflammation and remodeling. *Proc Natl Acad Sci* 2024;**121**:2017.
3. Dobin A, Davis CA, Schlesinger F, Drenkow J, Zaleski C, Jha S *et al.* STAR: ultrafast universal RNA-seq aligner. *Bioinformatics* 2013;**29**:15–21.
4. Liao Y, Smyth GK, Shi W. featureCounts: an efficient general purpose program for assigning sequence reads to genomic features. *Bioinformatics* 2014;**30**:923–930.
5. Love MI, Huber W, Anders S. Moderated estimation of fold change and dispersion for RNA-seq data with DESeq2. *Genome Biol* 2014;**15**:550.
6. Stephens M. False discovery rates: A new deal. *Biostatistics* 2017;**18**:275–294.
7. Liberzon A, Subramanian A, Pinchback R, Thorvaldsdóttir H, Tamayo P, Mesirov JP. Molecular signatures database (MSigDB) 3.0. *Bioinformatics* 2011;**27**:1739–1740.
8. Subramanian A, Tamayo P, Mootha VK, Mukherjee S, Ebert BL, Gillette MA *et al.* Gene set enrichment analysis: A knowledge-based approach for interpreting genome-wide expression profiles. *Proc Natl Acad Sci* 2005;**102**:15545–15550.
9. Liberzon A, Birger C, Thorvaldsdóttir H, Ghandi M, Mesirov JP, Tamayo P. The Molecular Signatures Database Hallmark Gene Set Collection. *Cell Syst* 2015;**1**:417–425.
