## Supplementary table 1 for "Small-molecule targeting MuRF1 protects against denervation-induced diaphragmatic dysfunction: Underlying molecular mechanisms"

Table S1.

| Source | Antibody | Dilution | Application |
| --- | --- | --- | --- |
| DSHB (BA-D5) | Anti-MyHC type I, mouse IgG2b | 1:800 | IF |
| DSHB (SC-71) | Anti-MyHC type IIa, mouse IgG1 | 1:800 | IF |
| Santa Cruz (sc-15376) | Anti-dystrophin, rabbit IgG | 1:500 | IF |
| Jackson ImmunoResearch (115-475-207) | DyLight goat anti-mouse IgG2b | 1:200 | IF |
| Jackson ImmunoResearch (115-225-205) | Cy2 goat anti-mouse IgG1 | 1:200 | IF |
| Jackson ImmunoResearch (711-165-152) | Cy3 donkey anti-rabbit IgG | 1:200 | IF |
| Cell Signaling (12293) | anti-SERCA1 | 1:1000 | WB |
| Abcam (ab83531) | anti-SERCA2 | 1:1000 | WB |
| Cell Signaling (14562) | anti-Phospholamban | 1:1000 | WB |
| Cell Signaling (5066) | anti-S100A1 | 1:1000 | WB |
| Cell Signaling (9101) | anti-phospho ERK1/2 (Thr202/Tyr204) | 1:1000 | WB |
| Cell Signaling (4695) | anti-ERK 1/2 | 1:1000 | WB |
| Cell Signaling (4058) | anti-phospho Akt (Ser473) | 1:1000 | WB |
| Cell Signaling (9272) | anti-Akt | 1:1000 | WB |
| Cell Signaling (2971) | anti-phospho mTOR (Ser2448) | 1:1000 | WB |
| Cell Signaling (2972) | anti-mTOR | 1:1000 | WB |
| Cell Signaling (9459) | anti-phospho 4E-BP1 (Thr37/46 ) | 1:1000 | WB |
| Cell Signaling (9644) | anti-4E-BP1 | 1:1000 | WB |
| Cell Signaling (9205) | anti-phospho P70S6K (Thr389) | 1:1000 | WB |
| Cell Signaling (9202) | anti-P70S6K | 1:1000 | WB |
| Myomedix (11005) | Anti-MuRF1 | 1:1000 | WB |
| Myomedix (8062) | Anti-MuRF2 | 1:1000 | WB |
| ECM (AP2401) | anti-Atrogin 1 | 1:1000 | WB |
| Cell Signaling (8081) | anti-K48 ubiquitinated protein | 1:1000 | WB |
| Abcam (ab22554) | anti-adiponectin | 1:1000 | WB |
| Cell Signaling (2443) | anti-PPARγ | 1:1000 | WB |
| Abcam (ab3527) | anti-perilipin1 | 1:1000 | WB |
| Cell Signaling (2125) | anti-alpha tubulin | 1:1000 | WB |
| Cell Signaling (7074) | anti-rabbit IgG HRP-linked antibody | 1:5000-10000 | WB |
| ECM (MS3001) | anti-mouse IgG HRP-linked antibody | 1:5000-10000 | WB |
| ThermoFisher (31402) | anti-goat IgG HRP-linked antibody | 1:5000-10000 | WB |
| Cell Signaling (7054) | anti-rabbit IgG AP-linked antibody | 1:1000 | WB |
