## Supplementary table 2 for "Small-molecule targeting MuRF1 protects against denervation-induced diaphragmatic dysfunction: Underlying molecular mechanisms"

Table S2.

| Sample ID | Experimental ID | Date | Conc. ng/uL NanoDrop | Conc. ng/uL Qubit RNA | A260 | A280 | 260/280 | 260/230 | Sample Type | Factor | RIN |
| --- | --- | --- | --- | --- | --- | --- | --- | --- | --- | --- | --- |
| 1 | Sham_2 | 2024/08/14 | 234.5 | 140 | 5.862 | 3.082 | 1.9 | 2.04 | RNA | 40 | 9.0 |
| 2 | Sham_3 | 2024/08/14 | 42.8 | 29.8 | 1.071 | 0.627 | 1.71 | 1.86 | RNA | 40 | 8.6 |
| 3 | Sham_4 | 2024/08/14 | 203.3 | 130 | 5.083 | 2.723 | 1.87 | 2.23 | RNA | 40 | 9.0 |
| 4 | Sham_5 | 2024/08/14 | 173 | 99.6 | 4.324 | 2.364 | 1.83 | 2.27 | RNA | 40 | 9.1 |
| 5 | Sham_6 | 2024/08/14 | 85.9 | 60.4 | 2.147 | 1.22 | 1.76 | 2.07 | RNA | 40 | 9.1 |
| 6 | Sham_7 | 2024/08/14 | 164.2 | 97.4 | 4.104 | 2.179 | 1.88 | 2.13 | RNA | 40 | 9.2 |
| 7 | Sham_8 | 2024/08/14 | 4.7 | too low | 0.118 | 0.062 | 1.91 | 1.23 | RNA | 40 | NA |
| 8 | DNV_VEH_2 | 2024/08/14 | 2.3 | too low | 0.058 | 0.033 | 1.78 | 0.12 | RNA | 40 | NA |
| 9 | DNV_VEH_3 | 2024/08/14 | 107.8 | 71.6 | 2.695 | 1.532 | 1.76 | 2.26 | RNA | 40 | 9.4 |
| 10 | DNV_VEH_4 | 2024/08/14 | 276.4 | 187 | 6.911 | 3.635 | 1.9 | 2.2 | RNA | 40 | 8.8 |
| 11 | DNV_VEH_5 | 2024/08/14 | 37.6 | 27.2 | 0.94 | 0.538 | 1.74 | 1.26 | RNA | 40 | 8.3 |
| 12 | DNV_VEH_6 | 2024/08/14 | 108.7 | 72.4 | 2.718 | 1.471 | 1.85 | 1.46 | RNA | 40 | 8.3 |
| 13 | DNV_VEH_7 | 2024/08/14 | 13.3 | 11.2 | 0.332 | 0.188 | 1.77 | 1.12 | RNA | 40 | NA |
| 14 | DNV_VEH_8 | 2024/08/14 | 65.5 | 48.4 | 1.638 | 0.917 | 1.79 | 1.75 | RNA | 40 | 8.7 |
| 15 | DNV_205_2 | 2024/08/14 | 6.2 | too low | 0.155 | 0.1 | 1.55 | 0.92 | RNA | 40 | NA |
| 16 | DNV_205_3 | 2024/08/14 | 6.5 | too low | 0.163 | 0.087 | 1.87 | 0.37 | RNA | 40 | NA |
| 17 | DNV_205_4 | 2024/08/14 | 16.6 | 36.6 | 0.416 | 0.228 | 1.82 | 1.18 | RNA | 40 | 8.0 |
| 18 | DNV_205_5 | 2024/08/14 | 560.1 | 338 | 14.003 | 7.307 | 1.92 | 2.28 | RNA | 40 | 8.3 |
| 19 | DNV_205_6 | 2024/08/14 | 388.7 | 394 | 9.718 | 4.989 | 1.95 | 2.13 | RNA | 40 | 8.7 |
| 20 | DNV_205_7 | 2024/08/14 | 37.7 | 40.6 | 0.942 | 0.53 | 1.78 | 1.82 | RNA | 40 | 8.6 |
| 21 | DNV_205_8 | 2024/08/14 | 49.8 | 41.6 | 1.245 | 0.743 | 1.68 | 2.08 | RNA | 40 | 8.2 |
