## Supplementary figures and images for "Small-molecule targeting MuRF1 protects against denervation-induced diaphragmatic dysfunction: Underlying molecular mechanisms"

### Supplementary figure 1

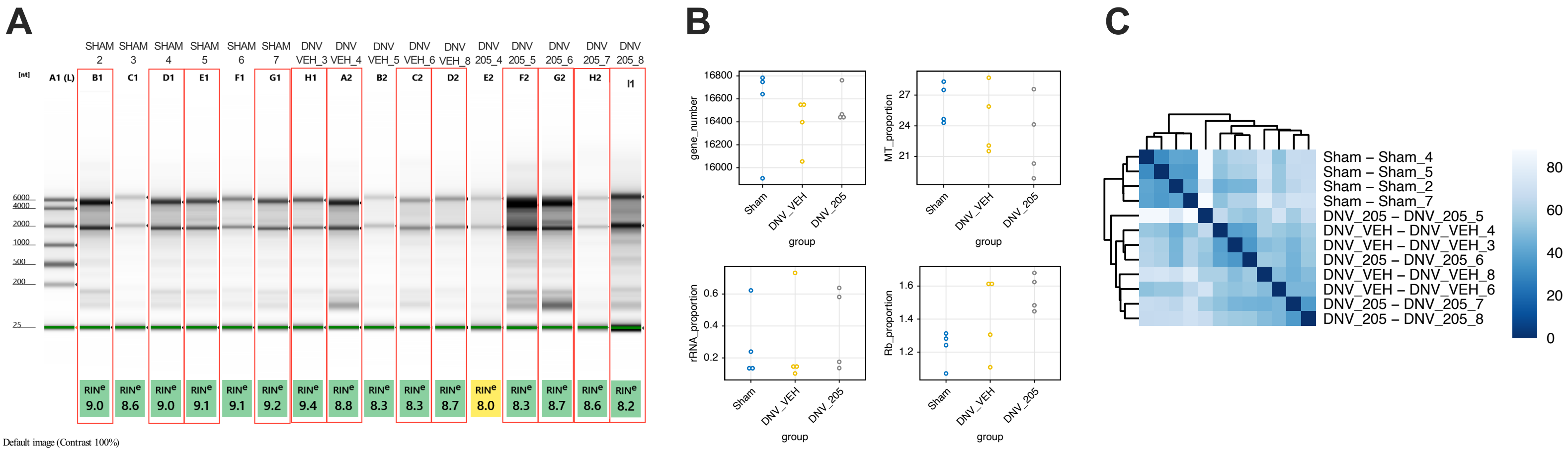

### Supplementary figure 2

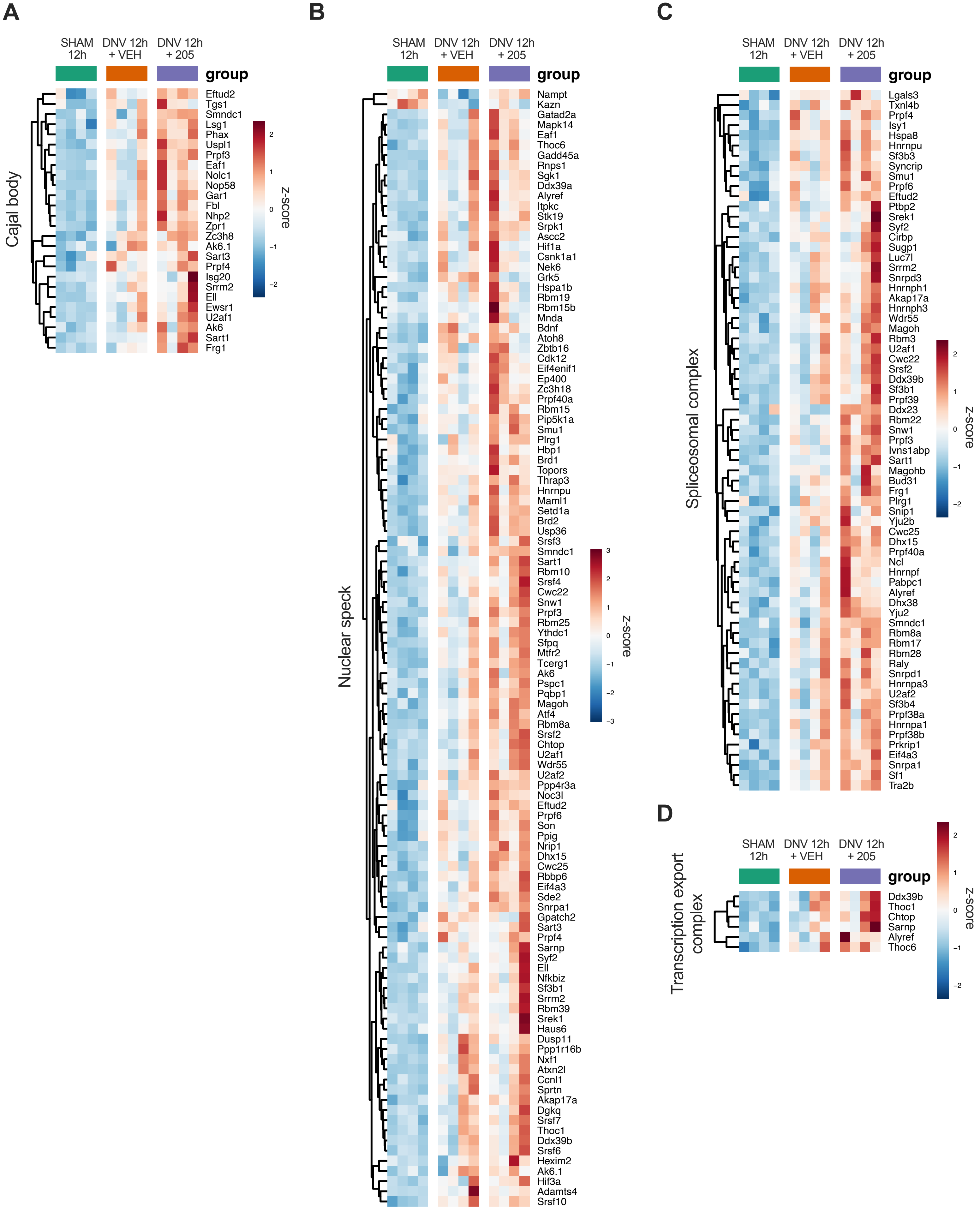

### Supplementary figure 3

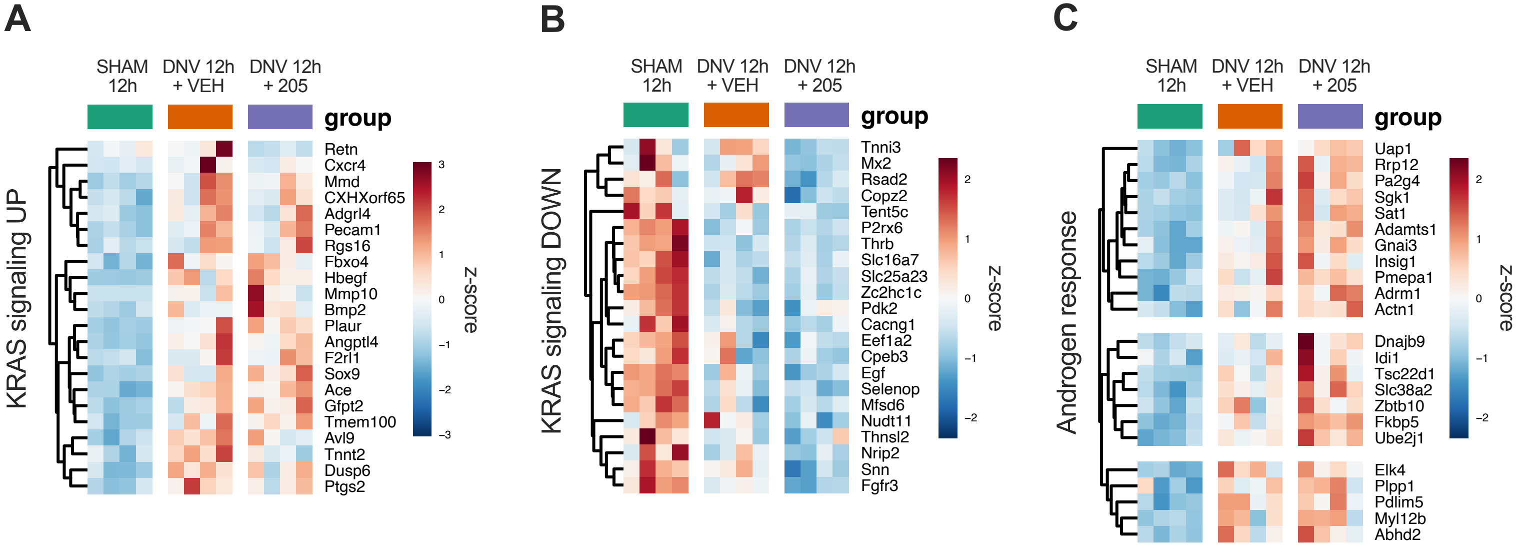

### Supplementary figure 4

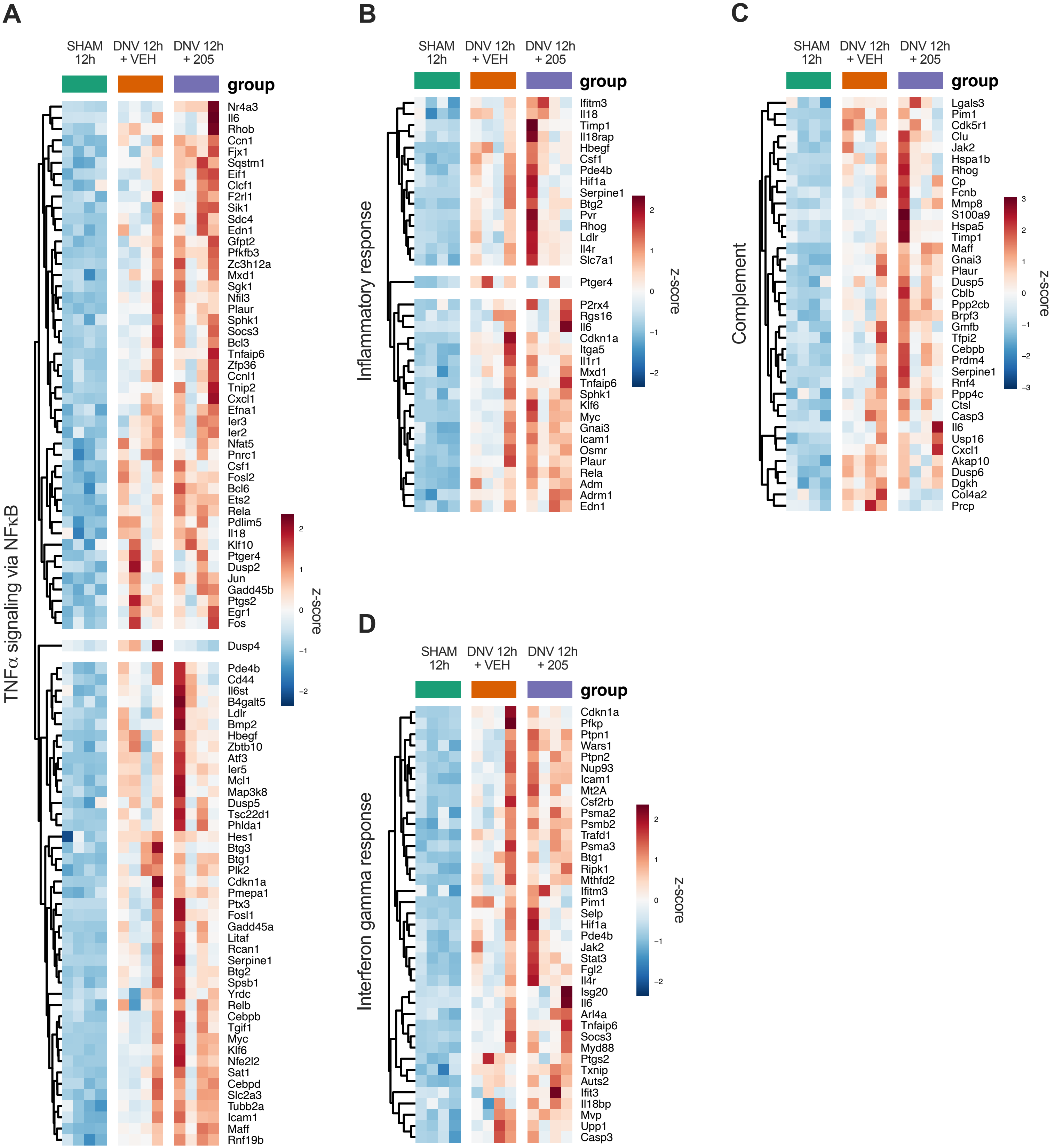

### Supplementary figure 5

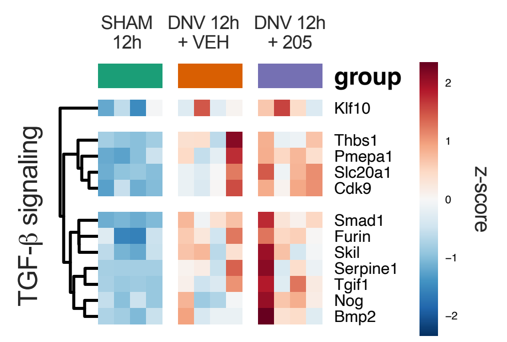
